# Longitudinal cancer evolution from single cells

**DOI:** 10.1101/2020.01.14.906453

**Authors:** Daniele Ramazzotti, Fabrizio Angaroni, Davide Maspero, Gianluca Ascolani, Isabella Castiglioni, Rocco Piazza, Marco Antoniotti, Alex Graudenzi

## Abstract

The rise of longitudinal single-cell sequencing experiments on patient-derived cell cultures, xenografts and organoids is opening new opportunities to track cancer evolution in single tumors and to investigate intra-tumor heterogeneity. This is particularly relevant when assessing the efficacy of therapies over time on the clonal composition of a tumor and in the identification of resistant subclones.

We here introduce LACE (Longitudinal Analysis of Cancer Evolution), the first algorithmic framework that processes single-cell somatic mutation profiles from cancer samples collected at different time points and in distinct experimental settings, to produce longitudinal models of cancer evolution. Our approach solves a Boolean matrix factorization problem with phylogenetic constraints, by maximizing a weighted likelihood function computed on multiple time points, and we show with simulations that it outperforms state-of-the-art methods for both bulk and single-cell sequencing data.

Remarkably, as the results are robust with respect to high levels of data-specific errors, LACE can be employed to process single-cell mutational profiles as generated by calling variants from the increasingly available scRNA-seq data, thus obviating the need of relying on rarer and more expensive genome sequencing experiments. This also allows to investigate the relation between genomic clonal evolution and phenotype at the single-cell level.

To illustrate the capabilities of LACE, we show its application to a longitudinal scRNA-seq dataset of patient-derived xenografts of BRAF^V600E/K^ mutant melanomas, in which we characterize the impact of concurrent BRAF/MEK-inhibition on clonal evolution, also by showing that distinct genetic clones reveal different sensitivity to the therapy. Furthermore, the analysis of a longitudinal dataset of breast cancer PDXs from targeted scDNA-sequencing experiments delivers a high-resolution characterization of intra-tumor heterogeneity, also allowing the detection of a late de novo subclone.

## Introduction

The advent of single-cell omics measurements has fueled an exceptional growth of high-resolution quantitative studies on complex biological phenomena [1, 2]. This is extremely relevant in the analysis of cancer evolution and in the characterization of intra-tumor heterogeneity (ITH), which is a major cause of drug resistance and relapse [3, 4, 5, 6].

In recent years, a large array of targeted cancer therapies has been developed, such as, e.g., kinase inhibitors, monoclonal antibodies and, more recently, immunomodulatory agents and clinical grade CAR-T [7]. However, the availability of such personalized therapies requires comparably advanced diagnostic and monitoring tools to study the response of cancer cells under the selective pressure generated by the treatment. In this respect, a highly-awaited major experimental advancement is provided by *longitudinal* single-cell sequencing experiments on samples taken at different time points from the same tumor, or from patient-derived cell cultures, xenografts or organoids [8, 9]. In most cases, single-cell transcriptomes are sequenced, e.g., via scRNA-seq experiments [10, 11], yet studies employing data from whole-genome/exome and targeted scDNA-seq experiments have been successfully proposed [12, 13]. Furthermore, approaches for the coupled analysis of genomes and transcriptomes at the single-cell level have been recently introduced [14, 15].

Longitudinal single-cell sequencing data might allow to track cancer evolution at unprecedented resolution, as they can be employed – in principle – to call somatic variants in each single cell of a tumor sample, at any given time point. Accordingly, this may allow to draw a high-resolution picture of evolutionary history of that tumor, as well as to measure the effect of any possible external intervention, such as a therapeutic strategy. Yet, currently no technique can explicitly process longitudinal single-cell mutational profiles.

On the one hand, in fact, the existing list of approaches that process single-cell data and extend phylogenetic methods by handling data-specific errors [16, 17, 18, 19, 20] are not directly applicable to multiple temporally ordered datasets, and cannot be used to investigate the clonal prevalence variation in time. On the other hand, even though methods for longitudinal *bulk* sequencing data are starting to produce noteworthy results [21, 22, 23], they usually require complex computational strategies to deconvolve the signal coming from intermixed cell subpopulations. Furthermore, there is an ongoing debate whether multi-sample trees from bulk samples are indeed phylogenies or, conversely, if they might lead to erroneous evolutionary inferences [24].

We here propose LACE (Longitudinal Analysis of Cancer Evolution), a new computational method for the reconstruction of longitudinal clonal trees of tumor evolution from longitudinal single-cell somatic mutation profiles of tumor samples.

In the inferred tree, each vertex correspond to a *genotype*, which identifies a subset of single cells displaying the same set of somatic variants; edges in the tree model both *parental* relations among genotypes and *persistence* relations through time. Notice that genotypes represent *(sub)clones* if the considered variants are *drivers*. Alternatively, if knowledge about drivers is not available, genotypes can be grouped into candidate subclones according to the co-occurrence similarity of mutations across single cells. LACE then estimates the prevalence of each candidate (sub)clone at each time point, hence allowing to identify (sub)clones that emerge, expand, shrink or disappear during the history of the tumor, e.g., as a consequence of a therapy or a selection sweep. A formal definition of the single-cell longitudinal clonal tree returned by LACE is provided in the Supplementary Information (SI).

Our method manages noise in single-cell sequencing data by estimating false positive and false negative rates – which might be different in distinct time points – and returns the longitudinal clonal tree that maximizes a weighted likelihood function computed on all data points. In this way, our method is able to handle possible differences in quality, sample size and error rates of the experiments performed at distinct time points. It is well known, in fact, that extremely different error rates are observed in single-cell experiments performed via distinct experimental platforms, and that even experiments made with the same platform might display highly heterogeneous noise levels [25]. The search is then performed by solving a Boolean matrix factorization problem [26], either via exhaustive search in case of very small models or via a Markov Chain Monte Carlo (MCMC), which ensures high scalability and convergence (with a large number of samplings).

The robustness of our approach allows its application to the highly-available scRNA-seq data – which are usually employed to characterize the gene expression patterns of single-cells in a variety of experimental settings [27] –, by calling somatic variants in transcribed regions with standard pipelines [28] and by selecting a set of confident variants, which might possibly include putative drivers. This allows to overcome the limitation of relying on longitudinal single-cell whole genome/exome sequencing experiments, which are currently rarer and significantly more expensive. In addition, by applying standard data analysis pipelines for the analysis of transcriptomes [27], our method allows to investigate the relation between somatic evolution and gene expression profiles in cancer clones and in single cells, especially in relation with possible external interventions, such as therapies.

There are several advantages in employing a formulation of the problem based on clonal trees, instead of standard phylogenetic trees in which single cells are placed as leaves. First, currently available single-cell data cannot guarantee the identifiability of a unique and reliable phylogenetic tree, mostly due to noise, to insufficient information and to the huge number of features of the output model [16]. This problem is significantly mitigated when considering clones instead of cells, as the model dimension is considerably reduced. Second, from the biological perspective, the resolution at the clone level is an effective choice to explain and predict cancer evolution and generate hypotheses with translational relevance [29], whereas phylogenetic trees including hundreds or even thousands of cells might be extremely difficult to query and interpret, especially with respect to the possible effect of therapies.

In order to assess the accuracy and robustness of the results produced by LACE, we performed extensive simulations, and compared with: SCITE [16] and TRaIT [19], two state-of-the-art tools for the inference of mutational trees from single-cell sequencing data, Sifit [18] a widely-used tool for tumor phylogenetic tree inference, SiCloneFit [20]) a Bayesian approach for the reconstruction of clonal trees from single-cell sequencing data, and with CALDER [23], a recent method for the reconstruction of longitudinal phylogenetic trees from bulk samples.

We applied LACE to longitudinal single-cell datasets of tumor samples generated with distinct experimental setups and, in particular, via either scRNA-seq or targeted scDNA-seq experiments. In the first case, we employed a longitudinal scRNA-seq dataset of patient-derived xenografts (PDXs) of BRAF^V600E/K^ mutant melanomas discussed in [30], by first performing mutational profiling via the widely-used GATK pipeline [31] and by selecting a panel of highly-confident somatic variants. In this respect, we here show that LACE can be used to produce robust results on longitudinal cancer evolution, even with noisy and incomplete data, and in particular, that it can characterize the efficacy of BRAF/MEK-inhibitor therapy on the clonal dynamics, also allowing to portray the phenotypic properties of the distinct (sub)clones. In the second case, we applied LACE to a longitudinal single-cell targeted DNA-seq dataset of PDXs generated from triple-negative breast tumors, presented in [32]. Our approach was applied on the set of somatic variants identified in the original work and allowed to refine the existing analysis, by producing a high-resolution picture of the evolutionary history of the tumor, thus proving the applicability of LACE to a wide range of existing data types.

## Results

### The LACE framework

LACE is a computational framework that processes multiple temporally ordered mutational profiles of single cells, collected from cancer samples or patient-derived cell cultures, xenografts or organoids, even in distinct experimental settings (e.g., pre- and post-treatment). Such profiles can be derived from whole-genome/exome or targeted scDNA-seq experiments, but also by calling variants from scRNA-seq data.

LACE takes as input a binary matrix for each time point/experiment, in which an entry is 1 if a somatic variant (e.g., single-nucleotide variants – SNVs, structural variants, etc.) is present in a given cell, 0 if not present and NA if the entry is missing. In order to select highly-confident variants, one can benefit from standard practices for variant-calling and from statistical filters. Furthermore, as one might be interested in selecting a set of putative drivers, which might possibly characterize (sub)clones, one can leverage on standard approaches for driver selection and on prior biological knowledge regarding the specific tumor type. Our method can be input with false positive and false negative rates for each time point, whenever such information can be derived from the technical features of the experiments. Otherwise, LACE includes a noise estimation procedure, performed via a parameter grid search.

LACE then solves a Boolean matrix factorization problem with standard phylogenetic constraints, by maximizing a weighted likelihood function on all time points. The rationale is that experiments collected at distinct time points may include even extremely different sample sizes and technical errors: LACE allows to balance such differences, by setting proper weights on the likelihood function. As default, the weights are set to be inversely proportional to the sample size of each dataset, in order to have comparable likelihood values through distinct experiments. LACE employs a MCMC search scheme on the phylogenetic matrix, which is defined by the association between genotypes/(sub)clones and sets of mutations, and which identifies a unique clonal tree. The weighted likelihood is maximized by exhaustively scanning the attachment of single cells to the genotypes/(sub)clones of the tree.

LACE returns:

- the maximum (weighted) likelihood clonal tree describing the longitudinal evolution of a tumor, in which nodes represent genotypes and edges are parental relations among them or persistence relations through time; a formal definition of the single-cell longitudinal tree returned by LACE is provided in the SI. Notice that genotypes represent *candidate (sub)clones* if the characterizing somatic variants are drivers.
- The attachment of single cells to genotypes/(sub)clones, which can be used to estimate the prevalence at any considered time point, as well as to identify macro evolutionary events, such as extinction or the emergence of new (sub)clones; as the cell attachment induces a partitioning of the single cells, this might be used in turn for mapping genotypes/(sub)clones on gene expression clusters, obtained via standard transcriptome analyses, if data are available.
- The error rates as estimated from data, when the grid search is employed.
- The corrected (theoretical) genotype of single cells.
- When no information on drivers is available, mutations can be grouped according to their similarity with respect to the corrected genotype matrix (see Methods). Intuitively, this should allow to identify which mutations mostly occur together in the cells of the considered dataset. Accordingly, the resulting groups of mutations can be used to identify candidate (sub)clones in the output model.

### Performance evaluation with synthetic simulations

In order to assess the performance of LACE and compare it with competing approaches, we performed extensive tests on synthetic datasets, generated with the cancer population dynamics simulator from [33] (the complete parameter settings of the simulations are provided as SI).

We generated a total of 900 independent datasets for distinct experimental scenarios (see below) and compared LACE with four state-of-the-art methods that process single-cell sequencing data from single time points, for the reconstruction of either mutational trees (SCITE [16] and TRaIT [19]), tumor phylogenetic trees (Sifit [18]), or clonal trees (SiCloneFit) [20]) –, and with a further method for phylogenetic tree inference from longitudinal bulk sequencing data (CALDER [23]). As synthetic datasets need to be adapted to be processed by such tools, with respect to CALDER, we computed the cancer cell fraction of driver mutations from the observed single-cell genotypes, and by assuming a uniform sampling of single cells and a read depth of 200X, which is typical for whole-exome sequencing experiments. Input data for SCITE, TRaIT, Sifit and SiCloneFit were generated by concatenating the longitudinal datasets in a unique mutational profile matrix.

We assessed the performance of methods by comparing: the reconstructed model with the ground-truth topology, in terms of precision, recall and accuracy; the inferred cell attachment matrix (i.e., the composition of clones) with the ground-truth, in terms of accuracy and adjusted Rand index; the corrected genotypes with the ground-truth, in terms of accuracy (please refer to the SI for a detailed description of all metrics).

### Synthetic longitudinal datasets generated from distinct experimental platforms

We designed a first in-silico scenario to account for sequencing experiments performed via distinct platforms at different time points, a likely setting for studies involving patient-derived cell cultures or organoids. In particular, we simulated the case in which the first and the third time points are characterized by a smaller number of cells 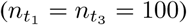 and a lower noise rate (false positive rate 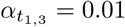, false negative rate 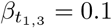), thus modeling a plausible setting resembling a Smart-seq protocol and the second time point in which a much larger number of cells is sequenced 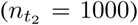, yet with a significantly higher noise rate 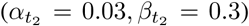, therefore modeling the features of a typical droplet-based experiment. We also assume that standard pipelines for mutation profiling from scRNA-seq data are employed, and we set a 0.5 probability for each dataset to have *γ* = 0.1 of missing entries.

In Fig. 2A one can see the distribution of precision and recall with respect to the ground-truth topology and of accuracy with respect to the ground-truth cell attachment matrix, for the six approaches in this scenario (the distribution of the tree accuracy, the precision-recall scatter-plots, the distribution of the adjusted Rand index for cell attachment matrix and the distribution of the corrected genotype accuracy, for this and the remaining experimental settings, are shown in the SI). By managing different error rates in distinct time points and by weighting the likelihood function with respect to the sample size, LACE is able to achieve the highest (median) values in both precision and recall with respect to the original topology and the best values of accuracy on cell attachment, showing in all cases the lowest variance, and outperforming all competing methods. This proves that LACE produces robust results when dealing with experiments from distinct protocols and with high differences in sample size and noise rates, as it might be common in real-world settings.

**Figure 1:**
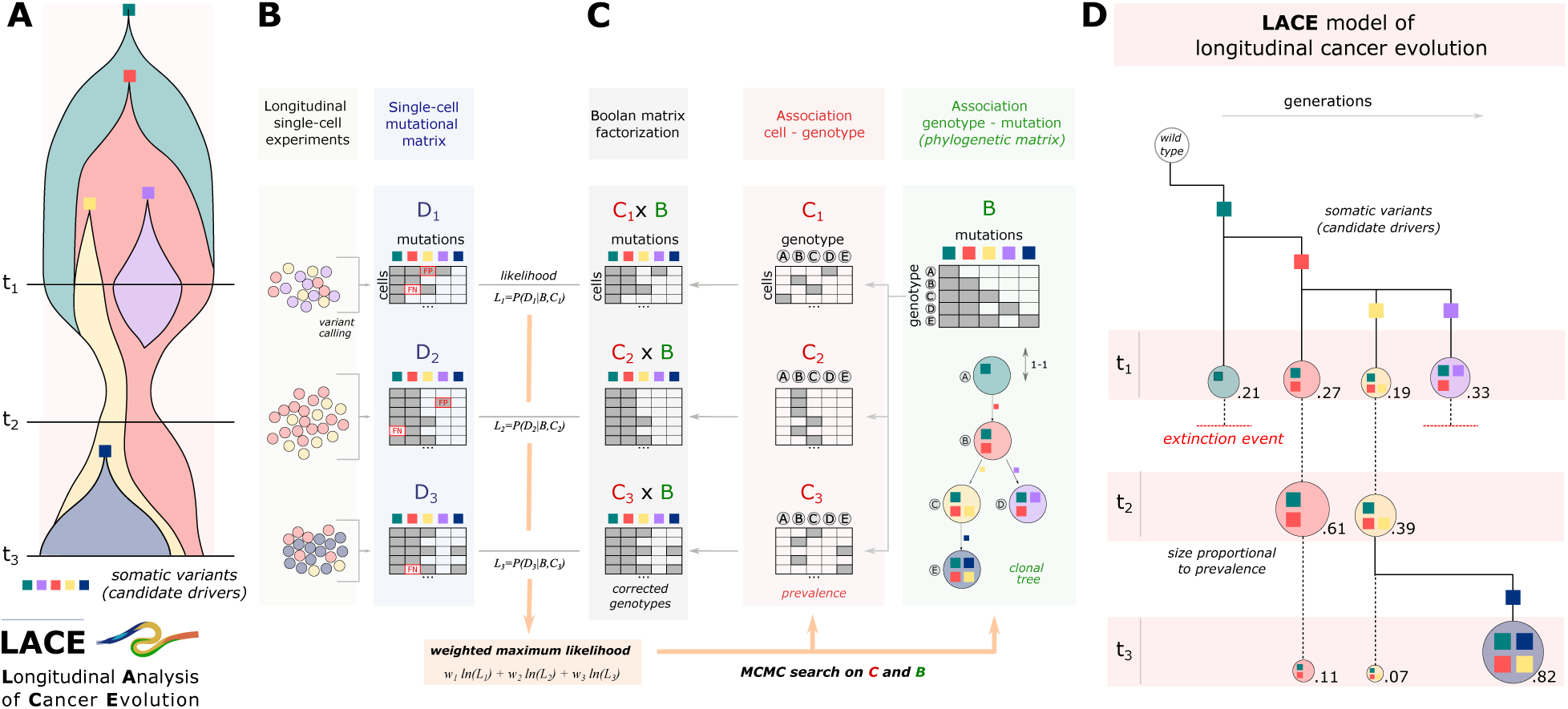
The LACE framework. **(A)** The branching evolution of a single tumor is described via a fishplot, and is characterized by the accumulation of 5 candidate driver mutations in distinct (sub)clones. **(B)** Single cells are sampled and sequenced at three subsequent time points: *t*_1_, *t*_2_ and *t*_3_, e.g., via scRNA-seq or (targeted) scDNA-seq experiments. By applying pipelines for variant calling, somatic mutation profiles for each time point are generated (matrices **D**_*i*_), which might include false positives, false negatives and missing entries. **(C)** LACE solves a Boolean matrix factorization problem **C**_*i*_ · **B** = **D**_*i*_, in which **B** is the phylogenetic matrix and **C**_*i*_ is the cell attachment matrix, by maximizing a weighted likelihood function computed on all time points, via a MCMC search scheme. **(D)** As a result, a unique longitudinal clonal tree is returned, which may include both observed and unobserved (sub)clones and the ancestral relations between them, as well as the prevalence variation, as measured at distinct time points. Solid lines represent parental relations between (sub)clones, each one characterized by a unique somatic variant (colored squares), whereas dashed lines represent persistence relations, which connect (sub)clones through time points or to extinction events.

**Figure 2:**
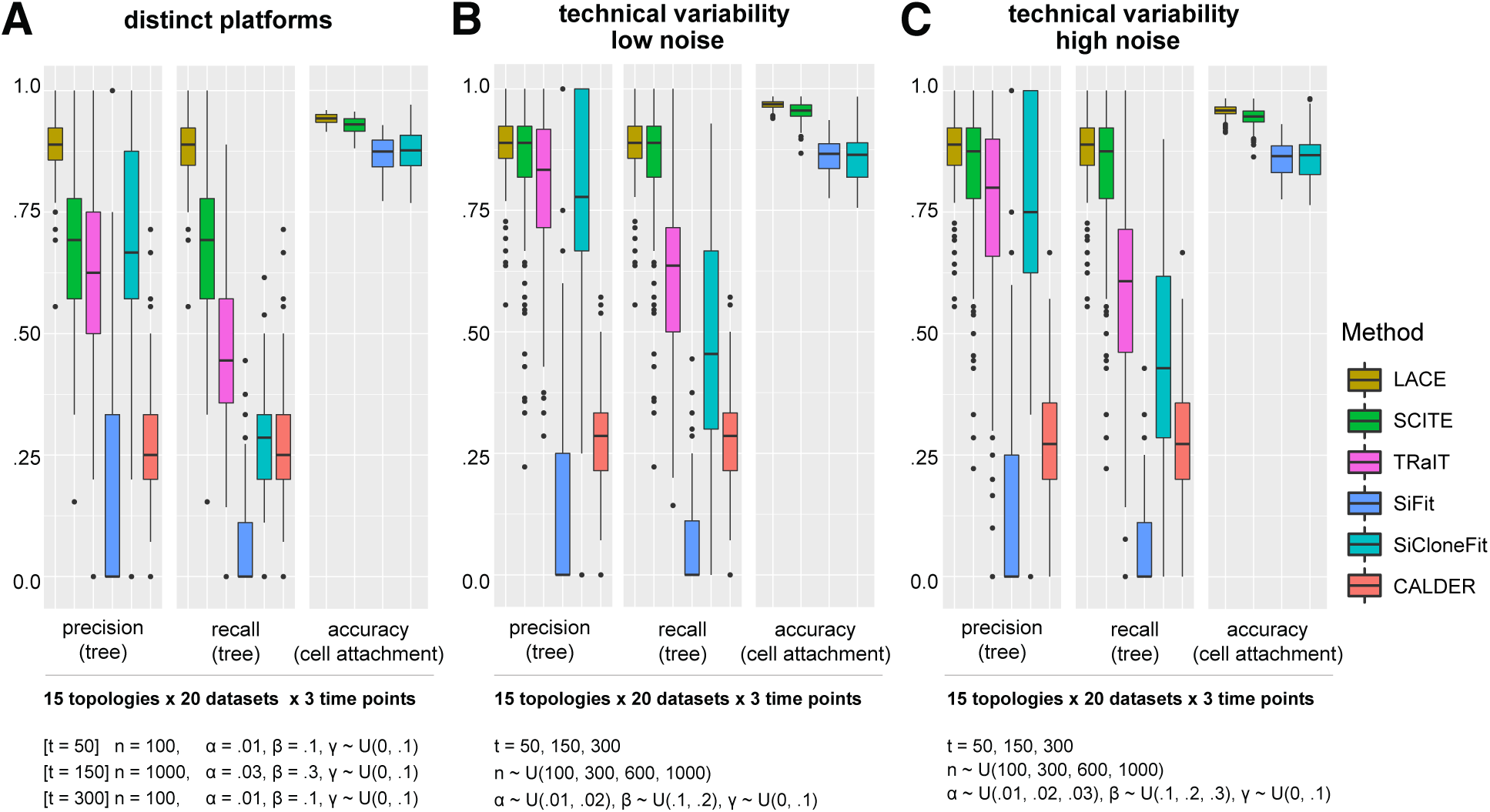
Comparison on simulated data. The population dynamics of cancer subpopulations was simulated with the tool from [34] – see the SI for the parameter setting. 15 scenarios were selected in which a number of drivers between 7 and 15 was observed during the simulation, resulting in branching evolution topologies. For each topology, a number of independent single-cell mutational profile datasets were sampled at 3 distinct time points of the dynamics. LACE was compared with SCITE [16], TRaIT [19], Sifit [18], SiCloneFit [20] and CALDER [23], on precision and recall with respect to the ground-truth topology and on accuracy with respect to the ground-truth cell attachment matrix (distributions are shown as boxplots; further metrics are provided in Supplementary Fig. 3). Notice that the accuracy on cell attachment cannot be computed for TRaIT and CALDER, as such methods do not return cell attachments or corrected genotypes. **(A)** In order to simulate single-cell data from distinct experimental platforms, we generated 20 independent datasets for each topology, including *n* = 100, *α* = 0.01, *β* = 0.1 for time points *t* = 50 and *t* = 300, and *n* = 1000, *α* = 0.03, *β* = 0.3 for time point *t* = 150. Each generated dataset was subsequently inflated with 10% of missing data, with a 0.5 probability. **(B)** In order to model technical variability, 20 independent datasets were sampled for each topology, in which at each time point (*t* = 50, 150, 300), each dataset includes with uniform and independent probability: a number of cells in the set {100, 300, 600, 1000}, *α* in the set {0.01, 0.02} and *β* in the set {0.1, 0.2}. Also in this case, approximately half of the datasets include 10% of missing data. **(C)** To account for higher error rates, 20 independent datasets for each topology were generated as in (B), with values of *α* in the set {0.01, 0.02, 0.03} and *β* in the set {0.1, 0.2, 0.3}.

### Synthetic longitudinal datasets with technical variability

To assess the consequences of noise and of technical variability in a larger experimental setting, we generated 300 independent datasets with a number of cells chosen at each time point with uniform probability in the set *n* = {100, 300, 600, 1000}. We modeled a first setting with low noise (i.e., *α* and *β* randomly chosen in the set {0.01, 0.02} and {0.1, 0.2}, respectively) and a second setting with higher values of noise (i.e., *α* and *β* in the range {0.01, 0.02, 0.03} and {0.1, 0.2, 0.3}, respectively).

In Fig. 2B-C one can see that also in this case LACE performs better than all competing methods in precision, recall with respect to the ground-truth topology and in cell attachment accuracy and proves to be robust with increasing noise levels (results on further metrics are consistent and are shown in the SI).

All in all, these results show the applicability of LACE to experimental settings in which a high variability in sample size and error rates is observed across different temporally ordered single-cell sequencing experiments. We also recall that LACE’s output is more expressive than those of methods designed to process single-time point datasets, as it allows to quantify the clonal prevalence variation in time, as well as to estimate the temporal positioning of phenomena such as clone emergence or extinction, for instance as a consequence of a therapy.

### Further experiments on simulated data

We further tested the robustness of LACE in a variety of simulated experimental scenarios. In particular, we assessed the inference accuracy when the real error rates are not provided as input and the grid search is employed. In Supplementary Fig. 6 one can see that LACE is robust and produces reliable results when employing the grid search on false positive/negative rates (please refer to the SI for details on the settings of this and the following experiments). Analogously, the results are stable even when wrongly specified error rates are provided as input to LACE (Supplementary File 3).

We also tested an in-silico scenario that accounts for the occurrence of back mutations due to, e.g., loss of heterozygosity, and which implies violations of the Infinite Sites Assumption (ISA) [35]. In Supplementary Fig. 5 one can see that LACE achieves optimal performances also in presence of relatively high proportions of variants affected by back mutations.

### Application of LACE to longitudinal scRNA-seq dataset from PDXs of BRAF-mutant melanomas

We applied LACE to a longitudinal dataset from [30]. In the study, the authors analyze multiple omics data generated from both bulk and single-cell experiments, to investigate minimal residual disease (MRD) in patient-derived xenografts from BRAF-mutant melanomas. In particular, they expose PDXs to BRAF^V600E/K^ inhibitor (i.e., *dabrafenib*), either alone or in combination with a MEK inhibitor (i.e., *trametinib*), and they perform multiple sequencing experiments at different time points.

Despite finding de novo mutations in known oncogenes (e.g., MEK1 and NRAS) in resistant cells, the analyses of the copy number alteration profiles, performed via massively parallel sequencing of single-cell genomes, was not sufficient to effectively characterize the clonal architecture and evolution of the tumor, whose composition appear to be similar prior to and after the treatment. Conversely, by analyzing transcriptomic data from both bulk and single-cell RNA-seq experiments, the authors were able to identify four distinct cell subpopulations, characterized by specific transcriptional states (i.e., *neural crest stem cell* – NCSC, *invasive, pigmented* and *starved-like melanoma cell* – SMC), which are insensitive to treatment and eventually lead to relapse, whereas the remaining cell subpopulations get quickly extinct. Based on these findings, the authors hypothesize that that the co-emergence of drug-tolerant states within MRD is predominantly due to the phenotypic plasticity of melanoma cells, which results in transcriptional reprogramming.

We here aim at refining the analysis of the clonal evolution of the tumor, by employing single-cell mutational profiles, as generated by calling variants from scRNA-seq data. In particular, we employed the GATK Best Practices [31] to identify good-quality variants and we finally selected 6 somatic SNVs, by applying a number of filters based on statistical and biological significance, and which might be considered as candidate drivers for this tumor (see Methods). We finally executed LACE with 50000 MCMC iterations, 100 restarts and by employing the grid-search on error rates.

The LACE model shown in Fig. 3A-B reveals the presence of a clonal trunk including a nonsynonymous somatic mutation on ARPC2 – a known melanoma marker [39] –, and of two distinct subclones, with somatic mutations on PRAME and RPL5 as initiating events. In particular, PRAME is a melanoma-associated antigen and known prognostic and diagnostic marker, which was recently targeted for immunotherapy [40]. RPL5 is a candidate tumor suppressor gene for many tumor types and displays inactivating mutations or focal deletions in around 28% of melanomas, which usually result in somatic ribosome defects [41].

**Figure 3:**
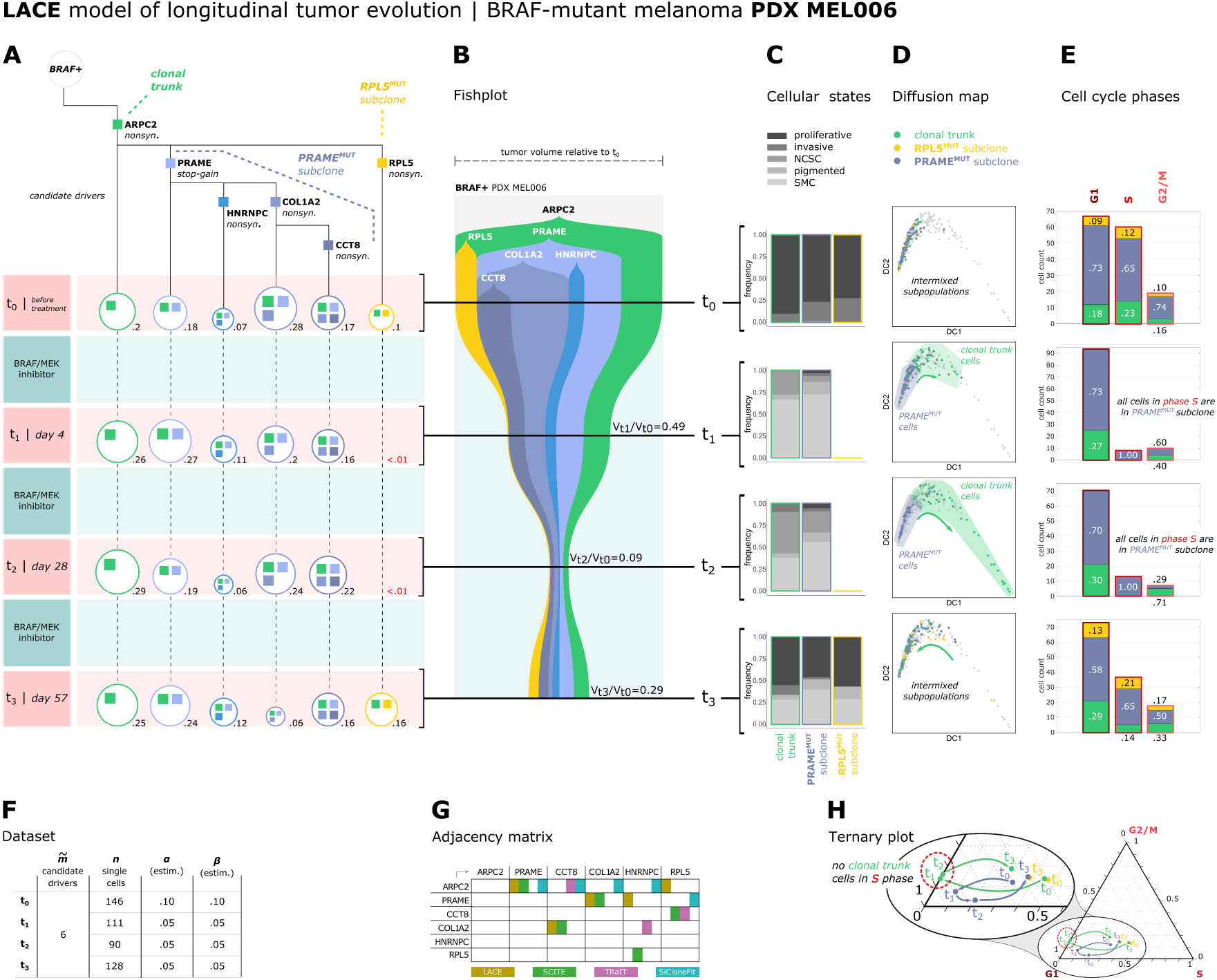
LACE model – PDX MEL006. **(A)** The longitudinal evolution of the PDX MEL006 derived from a BRAF mutant melanoma [30] returned by LACE is displayed. Single-cells were isolated and sequenced via scRNA-seq at four subsequent time points: (*t*_0_) before treatment (*n* = 146 single cells); (*t*_1_) after 4 days of concurrent treatment with BRAF inhibitor (i.e, dabrafenib) and MEK inhibitor (i.e, trametinib) (*n* = 111), (*t*_2_) after 28 days of treatment (*n* = 90), (*t*_3_) after 57 days of treatment (*n* = 128). Single-cell mutational profiles from scRNA-seq datasets were generated by applying the GATK Best Practices [31], and 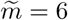 somatic variants were selected as candidate drivers to be used as input in LACE, with the procedure described in the main text. Each node in the output model represents a candidate (sub)clone, characterized by a set of somatic variants (colored squares). Solid edges represent parental relations, whereas dashed lines represent persistence relations. The clonal prevalence, as measured by normalizing the single-cell count, is displayed near the nodes and marked in red if lower than 1%. **(B)** The representation via a standard fishplot, generated via TimeScape [36], is displayed. The volume size at different time points is taken from [30]. **(C)** The composition of the clonal subpopulation (green) and of both the RPL5^MUT^ (yellow) and PRAME^MUT^ (blue) subclones with respect to the cellular states identified via single-cell transcriptomics analysis in [30] is shown for each time point (cells labeled by the authors as “other” were excluded from the analysis). **(D)** The diffusion maps [37] computed on 58 differentially expressed genes identified via ANOVA test (FDR adjusted *p <* 0.10) is shown; plots are generated via SCANPY [38]. A distinct diffusion map is shown for each time point, in which only the cells sampled at each time point are colored according to the clonal identity (i.e., clonal trunk, RPL5^MUT^ subclone or PRAME^MUT^ subclone). **(E)** The proportion of cells in G1, S and G2/M phases with respect to the distinct (sub)clones is shown with barplots for each time point. Cell phases are estimated on 97 cell cycle genes via SCANPY. At time point *t*_1_ and *t*_2_ all cells in phase S belong to the PRAME^MUT^ subclone. **(F)** In the table, the number of somatic variants 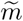, the number of single cells *n* and the estimated false positive and false negative rates, *α* and *β*, are returned for each time point. **(G)** The adjacency matrix describing the edges of the models returned by LACE, SCITE, TRaIT and SiCloneFit is shown. **(H)** A ternary plot showing the trajectories of the (sub)clonal subpopulations in the cell cycle space is shown. In the barycentric plot the three variables represent the ratio of cells belonging to phase G1, S and G2/M, respectively, and sum up to 1.

The PRAME^MUT^ subclone is characterized by the accumulation of further nonsynonymous mutations in HNRNPC, COL1A2 and CCT8 and displays an overall prevalence around ∼70% before treatment (*t*_0_). In particular, HNRNPC is a heterogeneous ribonucleoprotein involved in mRNA processing and stability, and its regulation appears to be involved in the PLK1-mediated P53 expression pathway, as it was shown via quantitative proteomic analysis on BRAF^V600/E^ mutant melanoma cells treated with PLK1-specific inhibitor [42]. COL1A2 is an extracellular matrix protein that is supposed to maintain tissue integrity and homeostasis, and was identified as a melanoma marker [43], as well as a candidate prognostic factors in several cancer types [44]. CCT8 encodes the theta subunit of the CCT chaperonin, and was found as up-regulated or mutated in several cancer types [45].

After 4 days of therapy (*t*_1_), the tumor volume halves. In particular, the RPL5^MUT^ subclone seems to disappear, whereas both the clonal subpopulation and the PRAME^MUT^ subclone maintain a stable prevalence. A further reduction of tumor volume is observed after 28 days of treatment (∼ 9% of the volume at *t*_0_), whereas at day 57 (*t*_3_) a significant growth is observed, in which the tumor reaches ∼ 29% of the initial volume, hinting at a possible relapse likely due to cells developing resistance. RPL5^MUT^ subclone reappears at time *t*_3_ with a prevalence around ∼ 16%, suggesting that its absence at time points *t*_1_ and *t*_2_ may be due to sampling limitations, which however do not affect the capability of LACE of inferring a correct evolution model. At time *t*_3_, the prevalence of all (sub)clonal subpopulations is similar to that of time *t*_0_, hinting at the absence of significant clonal selection. In Fig. 3G one can find the adjacency matrix representing the parental edges of the models inferred by LACE, SCITE, TRaIT and SiCloneFit (the complete models are shown in the SI). One can notice that, despite noteworthy differences, several phylogenetic relations, e.g., ARPC2 – PRAME, are retrieved by distinct methods.

We then analyzed the composition of (sub)clones with respect to the cellular states identified in [30] via single-cell transcriptomics analysis (Fig. 3C). Consistently with the findings in the article, the majority of cells in all (sub)clones are in a proliferative state before treatment, whereas at time point *t*_1_ and *t*_2_ no proliferative cells are left and all (sub)clones undergo transcriptional reprogramming, by displaying heterogeneous cell states (mostly starving-like melanoma cells, SMC), which result in acquired resistance at time *t*_3_, when cells restart proliferating in all (sub)clones.

As only minor differences are observed among (sub)clones with respect to cellular states, we refined the transcriptomic analysis, by first focusing on differentially expressed genes. The differential expression analysis, performed via standard ANOVA among all (sub)clones on all time points (data normalized by library size, FDR adjusted *p <* 0.10), allowed to identify PRAME as significantly up-regulated in PRAME^MUT^ cells, with log_2_-fold-change = 0.71; several other genes are found as differentially expressed among (sub)clones, yet with larger values of FDR p-value (the list of differentially expressed genes and the relative FDR p-values and fold-change values are provided in Supplementary File 1; please refer to the SI for further details on the analysis).

In order to analyze in depth the transition leading cells to resistance, we performed the same analysis with respect to the distinct time points. Interestingly, the list of differentially expressed genes among (sub)clones (FDR *p <* 0.10) includes no genes at time *t*_0_, *t*_1_ or *t*_3_, but it includes 58 genes at time *t*_2_ (see the Supplementary File 1). Within such group, 5 genes are significantly up-regulated in PRAME^MUT^ cells and display a log_2_-FC larger than 3, namely NGLY1 (log_2_-FC = 4.28), CDCA7 (log_2_-FC = 3.45), HK1 (log_2_-FC = 3.27), DNAJB4 (log_2_-FC = 3.27), ISOC2 (log_2_-FC = 3.11; the distribution of gene expression values at time *t*_2_ in PRAME^MUT^ and PRAME^WT^ cell subpopulations is shown in Supplementary Fig. 11). The results of this analysis suggest that, in addition to shifting their cellular states, distinct (sub)clones may differently respond to the therapy, and this would result in a transient increase of phenotypic heterogeneity, especially at time *t*_2_.

This aspect is particularly evident by looking at the projection of single cells in the space of the 58 most significant differentially expressed genes (FDR *p <* 0.10), represented via diffusion maps [37] in Fig. 3D. Before treatment (*t*_0_), almost all cells are positioned in the left region of the map and appear to be highly intermixed, proving the existence of a homogeneous phenotypic behaviour. At time *t*_1_, the RPL5^MUT^ subclone (yellow) disappears, whereas the clonal subpopulation (green) undertakes an apparent shift toward the right region of the map, which is characterized by transcriptional patterns that progressively diverge from those observed prior to the therapy, and which may possibly indicate high levels of cellular stress. This effect is notably amplified at time *t*_2_, where an explicit split of the clonal and the PRAME^MUT^ subpopulations can be observed, also in correspondence of the maximum dispersion of the cells on the map.

This outcome would further prove that distinct genetic clones may indeed suffer the effects of BRAF/MEK inhibition in different ways, during the resistance development phase, and this would result in different transcriptional patterns. At time point *t*_3_, when cells have achieved resistance and restart proliferating, RPL5^MUT^ subclone expands, and all cells appear to be intermixed on the left portion of the diffusion map once again.

We analyzed the cell cycle phase of the single cells at different time points, as estimated on 97 cell cycle genes via SCANPY [38]. In Fig. 3E one can see that cells are distributed across phases G1, S and G2/M in expected proportions in all subclones before therapy (*t*_0_). Strikingly, at time point *t*_1_ and *t*_2_ all cells in phase S belong to subclone PRAME^MUT^, whereas all cells of the clonal subpopulation are found in phase G1 or G2/M. At time *t*_3_ the scenario resembles that of time *t*_0_ and all (sub)clones include cells in all cell cycle phases. By looking at the ternary plot representing the proportion of cells in different cell cycle phases (Fig. 3H), it is possible to notice that the clonal and the PRAME^MUT^ subpopulations indeed undertake distinct trajectories in presence of therapy, before returning to a a state similar to the initial one.

This major result proves that the concurrent BRAF/MEK inhibition indeed affects in distinct ways different genetic clones during the resistance development stage. Apparently, cells lacking the PRAME mutation would be prevented from proceeding into S phase, whereas this effect would be highly mitigated in PRAME^MUT^ cells. All in all, these results cast a new light on the relation between clonal evolution and phenotype at the single-cell level, and suggest that distinct genetic clones may respond to therapy in significantly different ways.

In the SI we present a detailed analysis of the stability of LACE’s results for this case of study. In particular, we show that the consistency of the output model is preserved when either subsets or supersets of the selected somatic variants are considered and, similarly, when performing downsampling and oversampling of the original dataset (see Supplementary Figs. 13 and 14).

### Application of LACE to longitudinal targeted scDNA-seq dataset from PDXs of triple-negative breast tumors

We applied LACE to another longitudinal dataset from single-cell targeted DNA-sequencing experiments, presented in [32]. In the work, primary and metastatic tissues of breast cancer patients were serially transplanted into immunodeficient mice to generate xenograft lines, and the clonal dynamics was investigated by tracking single nucleotide and structural variants from bulk sequencing experiments. In addition, the authors performed targeted deep SNV re-sequencing to validate the clonal expansion of two xenograft series at the single-cell resolution. Sample SA501, in particular, was generated from a triple-negative breast primary tumor and was selected by the authors because of the interesting post-engraftment clonal dynamics.

We applied LACE by processing the allelic frequency matrix including a panel of 10 germline and 45 somatic variants selected by the authors, on the three engraftment passages for which single-cell deep re-sequencing was performed (i.e., X1, X2 and X4). In particular, we excluded germline variants and defined a binary mutational profile, by considering each somatic SNV in each single cell as: (*i*) present (1) if the allelic frequency *≥*0.10, (*ii*) absent (0) if the allelic frequency is *≤*0.01, (*iii*) missing entry (NA) if the allelic frequency is *>* 0.01 and *<* 0.10 or if already marked as uninformative (*<* 25 mapped reads). Finally, we filtered out all the variants displaying a value of missing data (NA) *≥*0.15 in all time points, to focus on highly confident SNVs. As a result, the mutational profile matrix provided as input to LACE includes 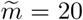 somatic variants and *n* = 27, 36, 27 single cells for time points *t*_0_ = X1, *t*_1_ = X2 and *t*_2_ = X4, respectively (Fig. 4B).

**Figure 4:**
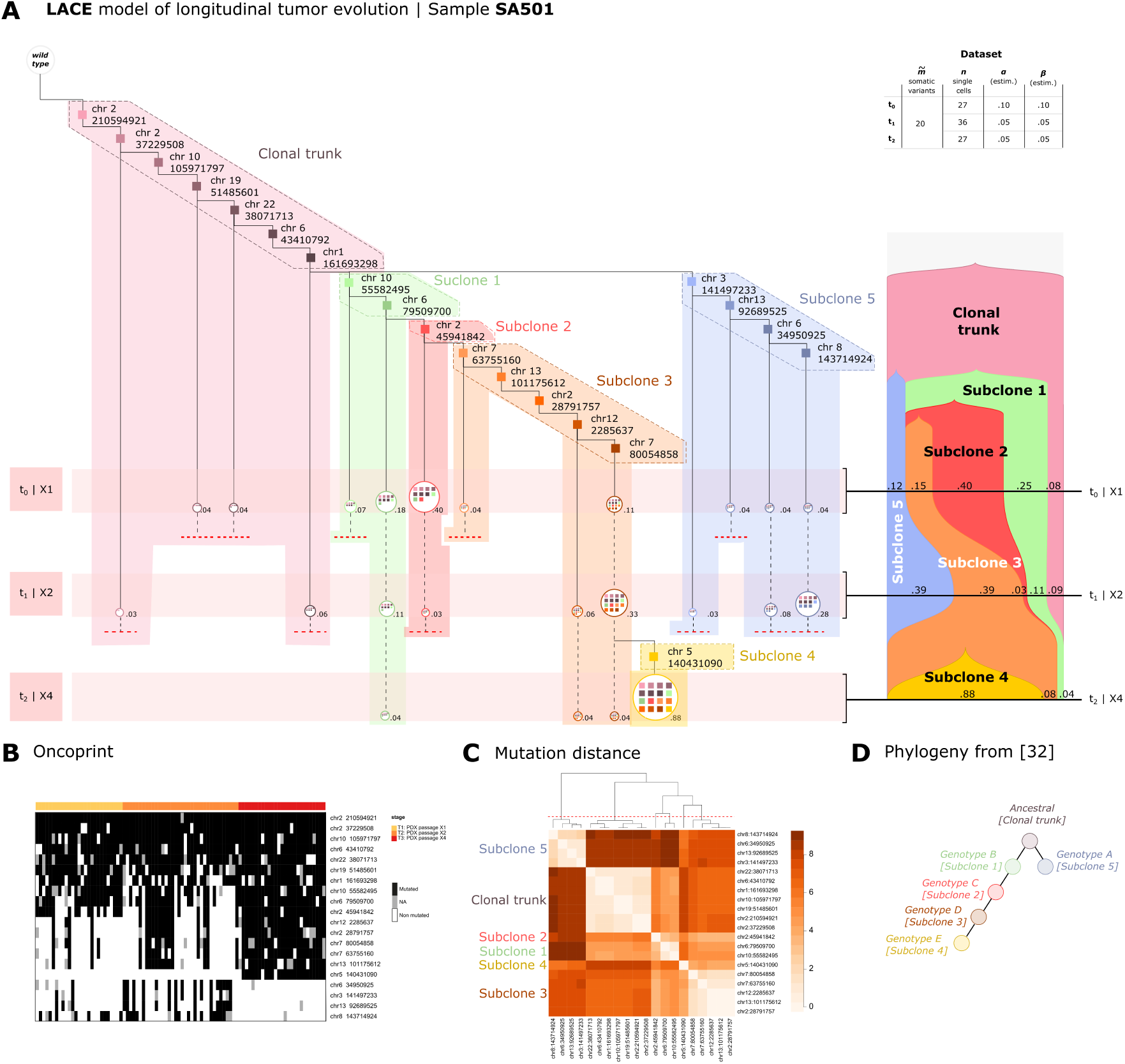
LACE model – Sample SA501. **(A)** The longitudinal evolution inferred by LACE on 3 PDXs generated from triple-negative breast tumor sample SA501 [32] is displayed. Targeted DNA sequencing experiments were performed on single cells of PDXs at three subsequent passages – *t*_0_ = X1 (*n* = 27 single cells), *t*_1_ = X2 (*n* = 36) and *t*_2_ = X4 (*n* = 27), by selecting a panel of 10 germline SNVs and 45 somatic SNVs. Single-cell mutational profiles were generated by discretizing the alternative allele ratio as proposed in the original work. Somatic SNVs displaying a ratio of missing data *>* 0.15 and germline mutations were filtered-out prior to the inference. As a result, 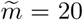 somatic variants were selected to be used as input in LACE. Each node in the output model represents a genotype characterized by a set of somatic mutations (colored squares). Solid edges represent parental relations, whereas dashed lines represent persistence relations. The prevalence of each genotype, as measured by normalizing the single-cell count, is displayed near the nodes. Colored shades mark 6 groups of mutations, which are identified by clustering variants according to the mutation distance defined in the text and shown in panel **C**, and which might represent candidate (sub)clones (the number of clusters was selected by following the analysis in [32]). In upper-right box, the number of somatic variants 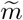, the number of single cells *n* and the estimated false positive and false negative rates, *α* and *β*, are returned for each time point. The representation via a standard fishplot, generated via TimeScape [36] at the resolution of candidate (sub)clones, is displayed on the right. **(B)** The oncoprint returning the single-cell mutational profile used in the analysis is shown. Black cells represent single cells in which the SNV is present, whereas gray cells represent missing values. **(C)** A heatmap returning the mutation distance among the 20 somatic variants used in the analysis is displayed (see Methods). 6 clusters identify candidate (sub)clones, and correspond to the 6 genotypes discussed in [32]. **(D)** The phylogenetic clonal model presented in [32] is displayed.

As no procedure for driver identification was employed in the original article, it may be sound to group the variants according to the patterns of co-occurrence across single cells, in order to identify candidate (sub)clones. In this respect, LACE allows to use a similarity measure based on the corrected genotype, which can be employed to cluster the variants of the output model (Fig. 4C; see Methods for further details). In Fig. 4A the LACE model of sample SA501 is displayed, which delivers a picture of the longitudinal evolution of the tumor at the resolution of both genotypes and candidate (sub)clones (LACE was executed with 50000 MCMC iterations, 100 restarts and by employing the grid-search on error rates).

6 candidate (sub)clones are identified by LACE, including a varying number of genotypes, and describing a complex clonal architecture. More in detail, a branching evolution scenario is observed, revealing the presence of two main subclones (#1 and #5), which appear from the clonal subpopulation prior to the first passage (*t*_0_). From subclone #1, further subclones emerge and are progressively selected through time, with a late subclone (#4) originating after time *t*_1_ and leading to a selection sweep at time *t*_2_ (clonal prevalence ∼88%). Interestingly, the evolution model returned by LACE is consistent with that proposed by the authors of the original work, who employed PyClone [46] to cluster mutations from bulk sequencing data and MrBayes [47] to reconstruct a phylogenetic model on single-cells (Fig. 4D). In particular, the cascading acquisition of mutations from parental to descendant clones observed by the authors is here refined by the explicit temporal ordering among genotypes provided by LACE, and which might used to generate hypotheses on driver identification (the cell attachment inferred by our analysis and that returned in the original work are provided in Supplementary File 2).

The application of LACE to a further dataset from [32] (sample SA494) is discussed in the SI (Supplementary Fig. 15), in which we show that our method is able to identify a rare clone at the metastatic level, prior to its expansion observed after transplanting in xenografts.

## Discussion

Cancer is an evolutionary process in which cells progressively accumulate somatic variants and undergo selection, while competing in a complex microenvironment [48]. Such variants can be used to track the evolution of a single tumor, to characterize intra-tumor heterogeneity and to identify the genomic makeup of subclones responsible for therapy resistance or phenotypic switches [49].

LACE is the first algorithmic framework that can process single-cell datasets collected at different time points to produce longitudinal clonal trees of tumor evolution. Remarkably, our approach can explicitly model different error rates and sample size in distinct experiments, which is typical in longitudinal studies. Accordingly, it can leverage the information extracted from possibly biased or non-exhaustive samplings of the tumor’s cells, which, instead, might lead to erroneous evolutionary inference when using single-time point datasets. Moreover, as the results are noise-tolerant, LACE can deliver reliable results even with extremely imperfect mutational profiles, as those derived by calling variants from transcriptome. This allows to exploit transcriptomic data, commonly available for most single-cell studies, to assess the clonal composition and the history of a tumor, and to directly investigate for the first time the relation between genomic clonal evolution and phenotype at the single-cell level, for instance in response to a certain treatment.

Importantly, new reliable protocols for high-quality genotyping of single cells from transcriptomic data are starting to appear, as proposed, for instance, with the Genotyping of Transcriptomes approach [15], in which the 10x Genomics platform is modified to amplify the targeted transcript and locus of interest. The improvement of the confidence on variant calling will accordingly enhance the accuracy of the results delivered by LACE. Furthermore, it was recently shown [50] that somatic mutations in mitochondrial DNA can be tracked by single-cell RNA sequencing and used for efficient lineage tracing. LACE might be easily employed with data on somatic mtDNA mutations from longitudinal cancer samples, once available. In any scenario, it is reasonable to expect a surge of high-quality single cell data on longitudinal cancer samples, which might be promptly used in LACE to track the evolution of tumors at unprecedented resolution.

Even though the application of longitudinal single-cell sequencing in clinical settings is still in its infancy, a rapid diffusion of these techniques is awaited, for instance in the context of hematological clonal disorders, where cancer cells are readily accessible over time. The availability of tools dedicated to the analysis of longitudinal single-cell data will allow the study of the detailed molecular mechanisms triggered by the therapies directly in primary cancer cells. Notably, in many hematological clonal disorders, such as Chronic Myeloid Leukemia, the leukemic stem cells and in particular the quiescent subset proved to be resistant even to targeted therapies such as Imatinib or second generation BCR-ABL1 inhibitors [51, 52]. Unfortunately, the nature of this resistance is still elusive, owing to the technical challenges involved in studying rare cell populations during treatment by using conventional approaches. In this scenario, LACE may be employed to characterize the resistance mechanisms selectively occurring in small subsets of the cancer cells pool.

As shown in the case studies, the innovative features of LACE allow to deliver experimental hypotheses with translational relevance. For instance, the model inferred from the BRAF-mutant melanoma PDX dataset produced a high-resolution picture of the evolutionary history of the tumor, as shaped by events such as the administration of a therapy. Furthermore, LACE allowed to identify (sub)clones that show different sensitivity to the therapy. In particular, we detected an unexpected behavior of the cell cycle machinery in different (sub)clones upon treatment, with only PRAME^MUT^ (sub)clone maintaining a fraction of cells in S phase. These findings suggest that longitudinal single-cell analyses are effective in dissecting the mechanisms by which cancer cells react upon treatment, at least for clonal disorders where tumor cells are readily accessible. All in all, by explicitly allowing a mapping between the clonal evolution and the phenotypic properties of single cells, LACE proved to be a powerful and expressive tool to decipher intra-tumor heterogeneity on multiple scales.

Our method was proven to be robust with respect to violations of the ISA on due to, e.g., back mutations; yet, the theoretical framework might be easily extended to account for possible violations of the ISA, as proposed, e.g., in [18, 53, 54], and by possibly leveraging the information on copy-number alterations. Furthermore, as both bulk and single-cell sequencing data may be increasingly available in longitudinal studies, the integration of both data types within our framework might allow to improve the clone identification and the inference quality, as proposed in [55, 29]. Finally, once a significant number of longitudinal single-cell datasets would be available on specific cancer types, techniques based on transfer learning might be applied to our models to identify possible patterns of recurrent evolution across tumors, as proposed by some of the authors in [56].

## Methods

### Input data preprocessing

LACE requires a distinct input data matrix including single-cell mutational profiles for any time point or experiment. As we are interested in somatic evolution of cancer subpopulations, mutational profiles may include, for instance, somatic single-nucleotide variants (SNVs) or structural variants.

Variant calling can be performed from DNA sequencing data – either whole-genome/exome or targeted sequencing – but also from transcriptomic data, such as scRNA-seq (see the SI for further details). The latter represents a cost-effective and highly-available alternative, despite known technological issues, such as the presence of reads encompassing intronic regions and the impossibility of calling variants from non-transcribed regions. In fact, LACE is robust with respect to high levels of noise, also thanks to the phylogenetic constraints implied by the process of accumulation of somatic mutations.

Notice that, as any statistical inference method, the reliability of the results is higher when the sampling bias is limited, as single-cell samplings should be ideally significant of the tumor’s composition at any time point. However, by exploiting the information extracted from multiple datasets sampled from the same evolutionary history, LACE can deliver statistically significant output models even with possibly biased, incomplete or imperfect samplings, as opposed to standard methods for single-time point data.

In our framework, a *genotype* defines a subset of single cells sharing the same set of somatic variants. In order to select high-quality variants, filters on statistical significance (e.g., recurrence thresholds) and on clinical/functional features should be employed, aimed at reducing the impact of noise of single-cell measurements and excluding non functional variants (e.g., rare polymorphisms).

Moreover, it may be sound to select somatic variants that might be candidate drivers for that specific tumor, i.e., mutations impacting the phenotypic variation among clonal populations. To this end, the presence of known variants involving oncogenes and tumor suppressor genes should be first verified. Previously uncharacterized mutations might be also selected, if significantly present in the samples. The identification of putative drivers allows to define *(sub)clones*, which we here consider as subsets of single cells sharing the same set of drivers. However, as driver identification is a typically hard task, at least when relying on data of single patients, one should prudently refer to *candidate* (sub)clones when interpreting the output model.

Furthermore, in case no reliable driver selection is possible, after the inference LACE allows to group genotypes according to the similarity of co-occurrence of mutations across single-cells (see below); in fact, groups with similar patterns of co-occurrence may indicate a candidate (sub)clone. In any case, the clonal evolution of the tumor inferred by our method is consistent at the resolution of both genotypes and (sub)clones, as LACE’s theoretical framework relies on the existence of a coherent process of accumulation of somatic variants.

### Single-cell data factorization problem (single time point)

Let us consider *k* different genotypes, *m* somatic variants and *n* single cells *c*_1_, …, *c*_*n*_ sampled in a given experiment. We can then define the following matrices:

1. The *single-cell data matrix* **D**: a binary *n × m* matrix where each row represents a single cell and each column a mutation; an element of **D**, *d*_*i,j*_ = 1 if we observe mutation *j* in cell *i*, otherwise *d*_*i,j*_ = 0.
2. The *phylogenetic matrix* **B**: a binary *k × m* matrix where each row represents a genotype and each column a mutation; *b*_*ij*_ = 1 if we observe mutation *j* in genotype *i*, otherwise *b*_*i,j*_ = 0. Each **B** can uniquely be represented by a tree and vice versa [57, 58]. Moreover, if we assume that the phylogenetic process is *perfect* and that the ISA holds [35], i.e., mutations are never lost and there is only one root. **B** has the following properties: (*i*) **B** is a square matrix (*k* = *m*), i.e., the number of genotypes is equal to the number of mutations; (*ii*) the rank of **B** is *k*; (*iii*) the Hamming distance between any pair of rows of **B** is *≥*1 and there is a column of **B** where all entries are 1; (*iv*) **B** is a lower triangular matrix and all the elements of its diagonal are equal to 1 [58].
3. The *cell attachment matrix* **C**: a binary *n × k* matrix, where each row represents a single cell and each column a genotype; *c*_*i,j*_ = 1 if cell *i* is associate to genotype *j*, otherwise *c*_*i,j*_ = 0. We notice that each cell is attached exactly to one genotype, i.e., the sum of any row of **C** is equal to 1. Note that this allows to easily compute the
4. prevalence of that genotype.

Notice that a genotype coincide with a (sub)clone when its characterizing somatic variants are drivers. On these premises and by assuming that the number of sampled cells is larger than the number of somatic variants, i.e., *n > m*, we can state the following proposition.

**Proposition 1** *For every single-cell matrix data* **D** *the following factorization holds:*

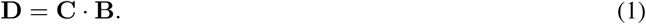

Its existence is guaranteed by construction, whereas the proof of uniqueness is straightforward.

### Likelihood function definition (single time point)

The factorization defined in Eq. (1) may not hold if **D** includes false positives, false negatives and missing values, which is typical in real-world scenarios. For this reason standard phylogenetic methods needs to be extended by modeling error rates, i.e., *α* as false positive rate and *β* as false negative rate, as proposed for instance in [16, 17].

By applying Bayes rule, the posterior probability of the output model (i.e., **B, C**) can be written as:

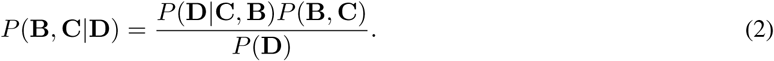

As typically done (e.g., in [16, 17]), we can safely remove *P* (**D**), which is an intractable integral and is identical for all models. Thus, we have:

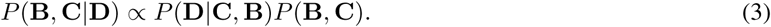

In the current implementation of LACE, we assume a uniform and uninformative prior on the phylogenetic matrix **B** and the cell attachment **C**. Whether knowledge on the underlying biological phenomenon would be available, our framework could be directly extended to include priors for *P* (**B, C**).

Let us define the estimated genotype matrix **G** = **C · B**, which is subsumed by the LACE model. Therefore, the problem is reduced to the maximization of the following likelihood function:

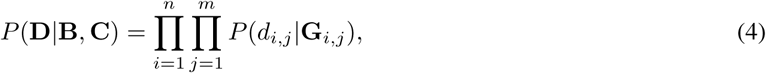

where *d*_*i,j*_ is a entry of **D** and:

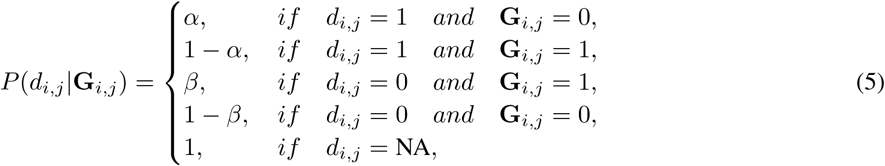

with *α* being the false positive rate, *β* the false negative rate, and NA labeling a missing entry. Notice that *α* and *β* can be provided as input to LACE, or they can be estimated (see below). We refer to the SI for details and discussions on how to account for high levels of noise in the input data by marginalizing the cell attachment.

### Longitudinal single-cell data factorization problem

We now generalize the problem to the case of *y* experiments taken at different time points *t*_1_ *≤ t*_2_ *≤ … ≤ t*_*y*_. For every experiment comprising *n*_1_, …, *n*_*y*_ different single cells and *m*_1_, …, *m*_*y*_ mutations, respectively, we can construct *y* independent data matrices **D**_1_, …, **D**_*y*_ as defined in (1).

As we assume that the longitudinal experiments are sampled from a unique generative phylogenetic matrix **B**, we need to expand the input matrices **D**_1_, …, **D**_*y*_ in order to include the union *m* of all the mutations detected in at least one cell over all the experiments. In detail, for each time point *s*, we define an expanded matrix 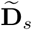, in which new columns corresponding to the mutations not detected at time point *s*, but detected in at least another time point, are added to **D**_*s*_. Each entry of such columns is then filled with 0 if the specific mutational locus has sufficient coverage (i.e., if the total read count exceeds a user-defined threshold) in that time point, otherwise it is set to NA. Notice that, by definition, 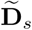 has 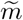 columns for *s* = 1, 2, …, *y*. Similarly, for each time point *s*, we define an expanded matrix 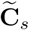, with 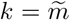 columns for *s* = 1, 2, …, *y*.

The longitudinal single-cell data factorization problem is then defined as follow:

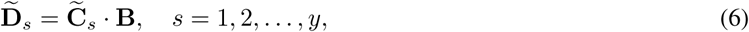

in which 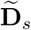 and 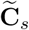 are the expanded matrices for *s*-th time point.

Notice that the formulation of an analogous problem for longitudinal bulk sequencing data was introduced in [23].

### Weighted likelihood function for longitudinal single-cell data

Since we are interested in learning a unique clonal tree **B** from different longitudinal datasets, we here define a weighted likelihood function as follows:

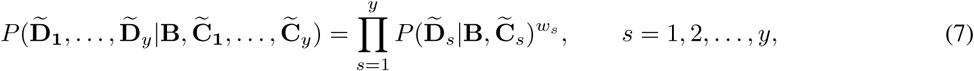

where 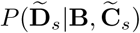 can be computed as for Eq. (4) and *w*_*s*_ are *weights* aimed at modelling possible idiosyncrasies of multiple longitudinal experiments, e.g., due to possible differences in quality and/or in the number of sampled cells. The definition of a weighted likelihood function allows us to explicitly account for experimental and technological differences among experiments collected at distinct time points and represents one of the major novelties of our approach. Note that in Eq. (7) we have *y* expanded estimated genotype matrices 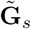

The choice of appropriate weights is problem- and data-specific and can benefit from a broad literature in statistical inference (see, e.g., [59]). In general, uniform weights would bias the solution toward datasets with larger sample sizes. For this reason, if no prior is available on the quality of the single experiments, we suggest as default weight for the *s*-th dataset composed by *n*_*s*_ cells: 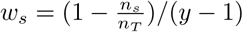, where *n*_*T*_ is the total number of cells of all the experiments, and *y ≥* 2 is the number of experiments.

Finally, we define the log-weighted likelihood objective function as follows:

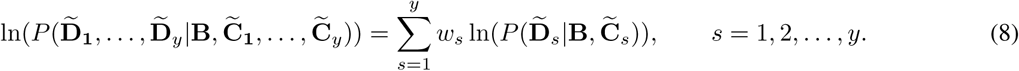

Notice that the values of *α* and *β* might be even extremely different among datasets – e.g., due to technological features of the experiments. LACE can explicitly model different error rates, which will be indicated as *α*_*s*_ and *β*_*s*_ for the *s*-th experiment.

By maximizing (8) LACE finally returns a corrected (theoretical) genotype matrix 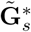 for each time point *s* = 1, …, *y*. The concatenation of such matrices can be named as 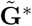 and will be employed in the definition of the mutation distance (see below).

### Error rates estimation

When error rates *α*_*s*_ and *β*_*s*_ are unknown, LACE includes a noise estimation procedure. By assuming that error rates are independent both from the phylogenetic matrix **B** and the cell attachments 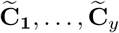, we can extend Eq. (7):

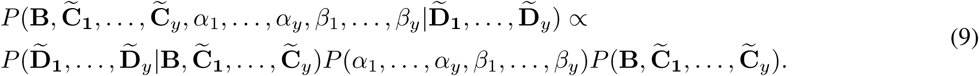

Finally, by assuming 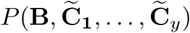 and *P* (*α*_1_, …, *α*_*y*_, *β*_1_, …, *β*_*y*_) to be uniform, our problem can be reduce to solving Eq. (8), given fixed values of *α* and *β*.

Therefore, the optimal values for *α*_1_, …, *α*_*y*_ and *β*_1_, …, *β*_*y*_ can be directly estimated by performing a parameter scan (see below).

### Search Scheme

LACE’s output model is composed by three components: the clonal tree **B**, the cell attachment matrices 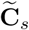, and (optionally) the estimated error rates *α*_*s*_ and *β*_*s*_, with *s* = 1, …, *y*, where *y* is the number of time points.

The search space of the possible solutions is huge, in fact given 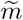 mutations (i.e., the union the mutations occurring at least once in all experiments) it includes continuous terms for *α*_*s*_ and *β*_*s*_, a discrete term of dimension 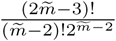 for **B**, and another discrete term of dimension 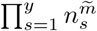 for 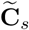.

As an exhaustive search can be achieved only for very small models, LACE employs a Markov Chain Monte Carlo (MCMC) scheme via a Metropolis–Hastings algorithm, to find the model that maximizes the weighted likelihood function defined in (8). In particular, the MCMC includes two ergodic moves on **B** : (*i*) node relabeling, (*ii*) prune and reattach of a single node and its descendants. For each proposed configuration **B**^*’*^, we find the maximum likelihood 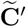 via an exhaustive search (the default probability for selecting move (*i*) or move (*ii*) is equal to 0.5).

Thus, given 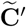 and by assuming that all the **B** are equally probable, the acceptance ratio *ρ* is given by:

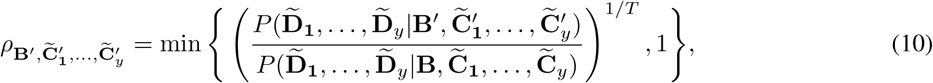

where *T* is a learning rate parameter, which could be used to speed up convergence as proposed in [16] (default value *T* = 1). With proper move probabilities, acceptance ratio and an infinite number of moves, the MCMC is ensured to converge to the maximum weighted likelihood solution [60, 16] (see the SI for tests on computation time and MCMC convergence). Notice that, as there might be multiple models with the same (maximum) weighted likelihood, in this case LACE returns all equivalent models as output.

In the current implementation of LACE we decided to exclude *α*_*s*_ and *β*_*s*_ from the MCMC, as they are continuous variables and this would dramatically increase the computational burden of the search. Therefore, LACE allows to perform a grid search on such parameters, by running multiple parallel MCMC searches with fixed error rates for each time point. The pseudocode of the algorithm can be found in the SI.

### Grouping genotypes via mutation distance

In case no driver identification strategy is employed in the selection of somatic variants, LACE allows to assess the presence of groups/clusters of mutations with similar co-occurrence patterns in the corrected genotype matrix, after the inference. Such groups might likely indicate (sub)clones and allow to provide a coarse-grained resolution to the evolution analysis (see, e.g., the analysis shown in Fig. 4).

Given the concatenation of the corrected (theoretical) genotype matrices 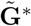 returned by LACE, the Euclidean distance between two variants *i* and *j* (i.e., columns of 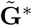), is given by:

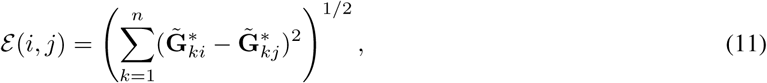

where *n* is the number of single cells in all experiments.

*ε* (*i, j*) can be then employed with standard clustering methods to identify groups of co-occurring mutations in the corrected genotype matrix 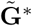 and indicating candidate (sub)clones.

### Synthetic data generation

To generate synthetic datasets we employed the tool from [34], which simulates a branching process modeling the population dynamics of cancer subpopulations, characterized by the accumulation of random mutations, which can either be drivers – i.e., inducing a certain proliferative advantage –, or passengers – i.e., with no effect. The simulator eventually returns the list of existing genotypes (including both passengers and drivers) and the relative prevalence at any time point, where a single time step corresponds to a generation (i.e., a replication event – see the SI for further details and the parameter settings).

We selected 15 simulation scenarios in which a number of drivers between 7 and 15 was observed, from which we sampled a large number of independent longitudinal single-cell mutational profile datasets including the drivers only, on three distinct time points, i.e., *t* = 50, 150 and 300. Single cells were randomly sampled with a probability proportional to cellular prevalence and we finally inflated the resulting binary mutational profiles with various rates of: false positives, *α*, false negatives, *β* and missing entries *γ*.

### Mutational profiling from scRNA-seq data

LACE requires longitudinal single-cell mutational profiles, as computed on a panel of selected somatic variants. Such profiles might be optimally derived by whole-genome and whole-exome single-cell sequencing experiments, which allow to call both known and uncharacterized somatic mutations, or by genotyping single cells, when target gene panels are available. In both cases, however, genome sequencing at the single-cell level is a still expensive and error-prone option, mostly related to technical issues in cell isolation and genome amplification [61].

Conversely, single-cell RNA-seq data are commonly used for an increasingly larger number of research goals and can be effectively used to call variants on exonic regions, even at the single-cell level [62]. It is known that scRNA-seq data are affected by different sources of noise [63] and cannot be used to call variants in non transcribed regions. However, we have shown that the results of LACE are noise-tolerant, also due to the phylogenetic constraints implied by the process of accumulation of mutations (see Fig. 2) and, therefore, it can be efficiently applied to possibly incomplete or noisy mutational profiles.

In particular, we here applied LACE to a longitudinal scRNA-seq dataset originally analyzed in [30]. In the study, a number of PDXs were derived from BRAF^V600E/K^ mutant melanoma patients and were treated with concurrent BRAF/MEK-inhibition. In our analysis, we selected PDX MEL006, for which four temporally-ordered scRNA-seq datasets are available: (*i*) pre-treatment, (*ii*) after 4 days of treatment, (*iii*) after 28 days of treatment, (*iv*) after 57 days of treatment. In the study, whole transcriptome amplification was made with a modified Smart-seq2 protocol and libraries preparation were performed using the Nextera XT Illumina kit. Samples were sequenced on the Illumina NextSeq 500 platform, by using 75bp single-end reads. Low-quality cells were filtered-out based on library size, number of genes expressed per cell, ERCCs, house-keeping gene expression and mitochondrial DNA reads, and a total of 674 cells was finally included in the dataset [30].

We applied further filters to remove cells displaying a fraction of counts on mitochondrial genes larger than 20% and cells displaying outlier values with respect to library size. As a result, we selected 475 single cells for downstream analysis.

In order to call SNVs and indels from such dataset, we employed the GATK Best Practices [31], which are proven to be effective even with single-cell data [62] (see the SI for a detailed description of the GATK pipeline). A VCF file including 272674 unique variants was generated and subsequently annotated with Annovar [64].

A first filtering step was applied to discard low-quality or non functional variants. First, synonymous and unknown mutations were removed (196320 unique mutations left); second, variants observed in less than two reads in each single cell were filtered-out (195931 unique mutations left); third, we employed a threshold of 1% on minor allele frequency, to remove possible germline mutations, as no normal tissue was included in the study (191599 unique mutations left).

A further filtering step was then employed to identify a list of putative drivers to be used in LACE. We first selected the mutations showing a frequency greater than 5% in at least one time point (595 unique mutations left). We then marked as missing entries (i.e., NA) the variants in a position with coverage lower than 3, as they might be miscalled due to gene expression down-regulation. We kept the mutations displaying less than 40% of missing data on all time points (151 unique mutations left), a median coverage larger than 10 and a median alternative read count larger than 4 (82 unique mutations left). As a final step, we manually curated the list of 82 remaining variants, to verify the possible presence of errors due to amplification (i.e., strand slippages) or alignment artifacts.

As a result, we selected the following 6 candidate drivers to be provided as input to LACE: ARPC2 (chr2:218249894, C>T, nonsynonymous substitution), CCT8 (chr21:29063389, G>A, nonsyn.), COL1A2 (chr7:94422978, C>A, non-syn.), HNRNPC (chr14:21211843, C>T, nonsyn.), PRAME (chr22:22551005, T>A, stop-gain), RPL5 (chr1:92837514, C>G, nonsyn.). The oncoprint including the mutational profiles of the single cells is displayed in Supplementary Fig. 7.

### Data availability

LACE is available as an open source R tool at: https://github.com/BIMIB-DISCo/LACE. The scRNA-seq dataset used in the first case study (PDX MEL006) was downloaded from GEO: https://www.ncbi.nlm.nih.gov/geo/, accession code: GSE116237. The targeted scDNA-seq datasets used in the second case study (PDXs SA501 and SA494) were retrieved from the Supplementary Material of the original article [32]. The source code used to replicate all our analyses, including synthetic and real datasets, is available at this link: https://github.com/BIMIB-DISCo/LACE-UTILITIES.

## Acknowledgements

This work was partially supported by the Elixir Italian Chapter and the SysBioNet project, a Ministero dell’Istruzione, dell’Università e della Ricerca initiative for the Italian Roadmap of European Strategy Forum on Research Infrastructures and by the AIRC-IG grant 22082. Support was also provided by the CRUK/AIRC Accelerator Award #22790, “Single-cell Cancer Evolution in the Clinic”. We thank Giulio Caravagna, Chiara Damiani, Francesco Craighero and Lucrezia Patruno for helpful discussions.

## Competing Interests

The authors declare that they have no competing financial interests.

## Contributions

D.R., F.A., D.M. and A.G. designed the approach, defined the method and implemented it. D.R. and A.G. performed the simulations. D.R., D.M., G.A., R.P. and A.G. executed the experimental data analysis pipeline. D.R., F.A., D.M., I.C., R.P., M.A. and A.G. analyzed the data and interpreted the results. A.G. and D.R. supervised the study. All authors discussed, drafted and approved the manuscript.

## Notes

https://github.com/BIMIB-DISCo/LACE

https://github.com/BIMIB-DISCo/LACE-UTILITIES

