## Supplementary Information for "Longitudinal cancer evolution from single cells"

### Supplementary Information: Longitudinal cancer evolution from single cells

**Daniele Ramazzotti[1,†], Fabrizio Angaroni[2,†], Davide Maspero[2,3,4,†], Gianluca Ascolani[2], Isabella Castiglioni[5,4], Rocco Piazza[1], Marco Antoniotti[2], Alex Graudenzi[4,*]**
[1] School of Medicine and Surgery, Univ. of Milan-Bicocca, Monza, Italy
[2] Dept. of Informatics, Systems and Communication, Univ. of Milan-Bicocca, Milan, Italy
[3] Fondazione IRCCS Istituto Nazionale dei Tumori, Milan, Italy
[4] Inst. of Molecular Bioimaging and Physiology, Consiglio Nazionale delle Ricerche (IBFM-CNR), Segrate, Milan, Italy
[5] Department of Physics "Giuseppe Occhialini", Univ. of Milan-Bicocca, Milan, Italy
† Equal contributors

### 1 Additional Materials and Methods

#### 1.1 Definition of single-cell longitudinal clonal tree

LACE returns as output: $\mathbf{B}$, i.e., the phylogenetic model, and $\tilde{\mathbf{C}}_i, \quad \forall i = 1, \dots y$, i.e., the expanded attachment matrix at time point $t_i$. The *single-cell longitudinal clonal tree* can be then defined as follows.

**Definition 1 (Single-cell Longitudinal Clonal Tree.)** *A single-cell longitudinal clonal tree is a node-weighted tree [1]: a triple* $\{V, E, P\}$, *where* $V$ *is the set of vertices* $E$ *is the set of the edges, and* $P$ *is the set of node weight functions* $p_k : V \to (0, 1), \quad p_k \in P$.

Indicating with $\tilde{c}^i_{j,l}$ the element at the $j$-th row and the $l$-th column of $\tilde{\mathbf{C}}_i$ and with $\mathbf{b}_z$ the $z$-th row of $\mathbf{B}$, it is possible to evaluate the sets $\{V, E, P\}$ as follows:

- **Vertices** : each element $v_h \in V$, $h = 1, \dots, r$ where $r$ is the total number of vertices in the model, is a couple $\{g, f\}$, where $g$ indicate the theoretical genotype associated to a vertex and $f$ the time point where it is observed.
  Given $\tilde{\mathbf{C}}_i$, for every time point we define a vector $\mathbf{a}^i = (a^i_1, \dots, a^i_k)^T$ where $k$ is the number of rows of $\mathbf{B}$ (i.e., the genotypes) and $a^i_j = \sum_{l=1}^{n} \tilde{c}^i_{j,l}$ (i.e., the sum of a column of the expanded attachment matrix $\tilde{\mathbf{C}}_i$). If $a^i_j \neq 0$, $g = \mathbf{b}_j$ and $f = t_i$. By repeating this procedure for all time points and rows of $\mathbf{B}$, one determine each element $v_h$ of the set $V$.
- **Edges**: the single-cell longitudinal clonal tree includes two different kinds of edges, i.e.:
  1. **Parental relation**: given two vertices $v_d = \{\mathbf{b}_r, t_a\}$, $v_e = \{\mathbf{b}_s, t_b\} \in V$, let $d(\mathbf{b}_r, \mathbf{b}_s)$ be the Hamming distance between the corrected genotypes associated to the vertices. $v_d$ is a parent of $v_e$, $v_d \xrightarrow{\text{PA}} v_e$ if:
  $$d(\mathbf{b}_r, \mathbf{b}_s) = 1, \quad \mathbf{b}_r \subset \mathbf{b}_s \quad \text{and} \quad t_a \leq t_b. \tag{1}$$
  Notice that this kind of edge represents the accumulation of a new mutation.
  2. **Persistence relation**: $v_d \xrightarrow{\text{PE}} v_e$ represents a persistence relation from vertex $v_d = \{\mathbf{b}_r, t_a\} \in V$ to vertex $v_e = \{\mathbf{b}_s, t_b\} \in V$, if:
  $$d(\mathbf{b}_r, \mathbf{b}_s) = 0 \quad \text{and} \quad b - a = 1. \tag{2}$$
  Notice that persistence relations are depicted via dashed edges in the output model.
- **Weights**: $P$ is a set of functions $p_k : V \to (0, 1)$ such that $p_k(v_k) = p_k(\{\mathbf{b}_r, t_i\}) = \sum_{l=1}^{n_i} \frac{\tilde{c}^i_{r,l}}{n_i}$, where $n_i$ is number the cells present in the time point $t_i$. Such weights represent the prevalence of the vertices at every time points.

#### 1.2 Marginalization of cell attachment

In case of extremely noisy data, it might be sound to marginalize the attachment of cells to clones (i.e., $\mathbf{C}$), as proposed in [2]. In our formalism, this is achieved by summing among all possible cell attachments. The likelihood becomes:

$$P(\mathbf{D}|\mathbf{B}) \propto \prod_{s=1}^{n} \sum_{i=1}^{k} \prod_{l=1}^{m} P(d_{s,l}|\mathbf{G}_{s,l}). \tag{3}$$

The marginalization cell attachments might be useful to speed up the convergence of the MCMC in the case of very noisy data.

#### 1.3 MCMC convergence test

In order to assess the convergence of the MCMC scheme, we executed the standard Geweke diagnostic test [3] on 100 independent computations performed on a selected synthetic dataset – Setting (A) distinct platforms (see Supplementary Table 1), $\tilde{m} = 15$ variants and $n = 1200$ single cells – with 1000 MCMC iterations and 10 restarts. The diagnostic tests the equality of the means of the first 10% and last 50% of the MCMC. In case the samples are drawn from a stationary distribution of the MCMC, the Geweke's statistic – computed as a Z-score – displays an asymptotically standard normal distribution (the Z-score tends toward 0). Otherwise, if the two parts of the chain are not drawn from the same distribution, it is sound to discard the first iterations to check if and when the rest of the chain has converged.

For this reason, we repeated the Geweke diagnostic test in 5 scenarios in which we discarded an increasingly larger number of iterations from the beginning of the chain. In detail, we divided the first 50% of the chain in 4 equal segments, and computed the Z-score between the first 10% and last 50% of the resulting chain, in the following 5 scenarios: $(i)$ with the whole chain ($1^{\text{st}}$ – $1000^{\text{th}}$

iteration), $(ii)$ by discarding the first 10% of the chain (100$^{\text{th}}$ – 1000$^{\text{th}}$ iteration), $(iii)$ by discarding the first 20% of the chain (200$^{\text{th}}$ – 1000$^{\text{th}}$ iteration), $(iv)$ by discarding the first 30% of the chain (300$^{\text{th}}$ – 1000$^{\text{th}}$ iteration), $(v)$ by discarding the first 40% of the chain (400$^{\text{th}}$ – 1000$^{\text{th}}$ iteration).

In Supplementary Fig. 1 one can see the distribution of Geweke's Z-scores in such scenarios, in relation to a benchmark line representing 2 standard deviations (dashed red line). One can notice that both scenarios $(i)$ and $(ii)$ display distributions that are significantly divergent from the standard normal assumption (Z-score $\sim 0$), thus showing that the burn-in of the chain is reached around 300 iterations and proving its convergence.

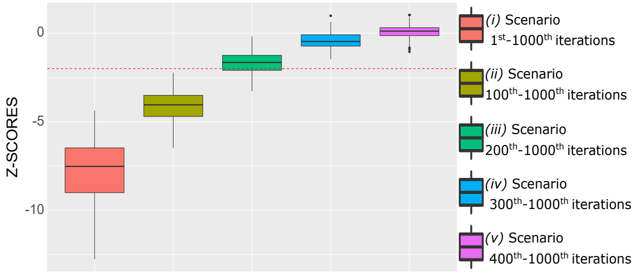

Figure 1: **Geweke diagnostic test.** Z-scores distribution from the Geweke diagnostic, with respect to the scenarios discussed in the test and characterized by progressively smaller chain intervals. The dashed red line represents 2 standard deviations from the normal standard assumption. We observe that the standard normal assumption falls around 300 iterations of the Markov chain.

#### 1.4 Computation time

We extensively assessed the computation time of LACE in different simulated scenarios, by especially evaluating the impact of variations in the number of variants $\tilde{m}$ and in the number of single cells $n$. All simulations were performed by employing the simulation setting A – distinct platforms, described in Supplementary Table 1.

In the first scenario, we assessed the variation of $\tilde{m}$, with respect to three distinct configurations: $(A-i)$ $\tilde{m}=5$ variants, $n_1=100$, $n_2=1000$, $n_3=100$ single cells; $(A-ii)$ $\tilde{m}=10$, $n_1=100$, $n_2=1000$, $n_3=100$; $(A-iii)$ $\tilde{m}=15$, $n_1=100$, $n_2=1000$, $n_3=100$. In the second scenario, we evaluated the variation of $n$, with respect to three further configurations: $(B-i)$ $\tilde{m}=15$, $n_1=10$, $n_2=100$, $n_3=10$; $(B-ii)$ $\tilde{m}=15$, $n_1=50$, $n_2=500$, $n_3=50$; $(B-iii)$ $\tilde{m}=15$, $n_1=100$, $n_2=1000$, $n_3=100$. For each configuration, we executed 100 independent computations, with 1000 MCMC iterations and 5 restarts, on a single core of a MacBook Pro, processor 2.6 GHz Intel Core i7, 16 GB RAM 2400 MHz DDR4. The distribution of the run times with respect to the distinct configurations is shown in Supplementary Fig. 2.

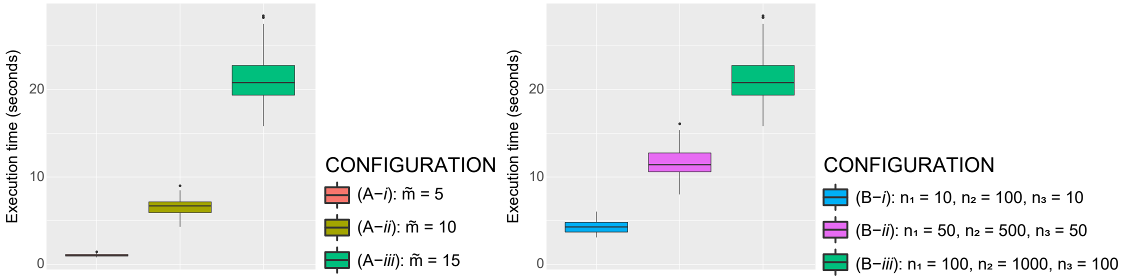

Figure 2: **Computation time**. Computation time of LACE with respect to the 6 different configurations described in the text; every boxplot shows the distribution of 100 independent computations with 1000 MCMC iterations and 5 restarts. Execution time is provided in seconds.

##### 1.5 Pseudocode

**Inputs:**
$\mathcal{D} = \{\tilde{\mathbf{D}}_1, \cdots, \tilde{\mathbf{D}}_y\}$, *expanded mutation data for each time point* $y$
$w = \{w_1, \cdots, w_y\}$, *likelihood weights for each time point* $y$
$\alpha = \{\alpha_1, \cdots, \alpha_y\}$, *false positive rates for each time point* $y$
$\beta = \{\beta_1, \cdots, \beta_y\}$, *false negative rates for each time point* $y$
$\lambda$, *number of MCMC restarts*
$\delta$, *maximum number of MCMC steps*
**Outputs:**
$\mathbf{B}$, *phylogenetic matrix*
$\mathcal{C} = \{\tilde{\mathbf{C}}_1, \cdots, \tilde{\mathbf{C}}_y\}$, *cell attachment matrices for each time point* $y$
$LL_{single}$, *log-likelihood for each time point* $y$
$LL_{weighted}$, *weighted log-likelihood*
$clonal\_prevalence$, *clonal prevalence*
$\mathbf{B} = \emptyset$
$\mathcal{C} = \emptyset$
$LL_{single} = \emptyset$
$LL_{weighted} = -\inf$
**for** $(i \in 1 : \lambda)$ **do**
  $\mathbf{B}_{curr}$ = **InitializeB(**$\mathcal{D}$**)**
  $\{\tilde{\mathbf{C}}_{curr}, LL_{single_{curr}}, LL_{weighted_{curr}}\}$ = **MaximumLikelihood(**$\mathbf{B}_{curr}$**,** $\tilde{\mathbf{D}}$**,** $w$**,** $\alpha$**,** $\beta$**)**
  **for** $(j \in 1 : \delta)$ **do**
    $\mathbf{B}_{tmp}$ = **MCMCMove(**$\mathbf{B}_{curr}$**)**
    $\{\tilde{\mathbf{C}}_{tmp}, LL_{single_{tmp}}, LL_{weighted_{tmp}}\mathbf{B}\}$ = **MaximumLikelihood(**$\mathbf{B}_{tmp}$**,** $\tilde{\mathbf{D}}$**,** $w$**,** $\alpha$**,** $\beta$**)**
    $rand = U \sim (0, 1)$
    **if** $(rand \leq min(exp(LL_{weighted_{tmp}} - LL_{weighted_{curr}}), 1))$ **then**
      $\mathbf{B}_{curr} = \mathbf{B}_{tmp}$
      $\tilde{\mathbf{C}}_{curr} = \tilde{\mathbf{C}}_{tmp}$
      $LL_{single_{curr}} = LL_{single_{tmp}}$
      $LL_{weighted_{curr}} = LL_{weighted_{tmp}}$
    **end**
  **end**
  **if** $(LL_{weighted_{curr}} > LL_{weighted})$ **then**
    $\mathbf{B} = \mathbf{B}_{curr}$
    $\tilde{\mathbf{C}} = \tilde{\mathbf{C}}_{curr}$
    $LL_{single} = LL_{single_{curr}}$
    $LL_{weighted} = LL_{weighted_{curr}}$
  **end**
**end**
$clonal\_prevalence$ = **ComputeClonalPrevalence(**$\mathbf{B}, \tilde{\mathbf{C}}$**)**

**Algorithm 1:** Pseudocode describing LACE algorithm.

##### 1.6 Performance assessment metrics

To compare the performance of LACE and competing methods over synthetic data, we evaluated the following metrics:

- **tree metrics** :
  1. *precision*: $\frac{\text{TP}}{\text{TP+FP}}$,
  2. *recall*: $\frac{\text{TP}}{\text{TP+FN}}$,
  3. *accuracy*: $\frac{\text{TP+TN}}{\text{TP+TN+FP+FN}}$,

  where TP are the true positives, (i.e., rightly inferred edges), FP the false positives (i.e., wrongly inferred edges), FN the false negatives (i.e., edges that are present in the ground-truth topology, but are not inferred), TN the true negatives (i.e., edges that are not inferred and are not present in the ground-truth topology).
- **genotype metrics**:

1. *accuracy on cell attachment* (see above),
   as computed with respect to the ground-truth cell attachment matrix $\tilde{C}_s \quad s = 1, \ldots, y$. In this case true positives are the cells correctly associated to a genotype, whereas false positives are cells wrongly associated to a genotype;
2. *adjusted Rand index on cell attachment*: ARI $= \frac{\sum_{ij}\binom{q_{ij}}{2} - \left[\sum_i \binom{a_i}{2} \sum_j \binom{b_j}{2}\right] / \binom{q}{2}}{\frac{1}{2}\left[\sum_i \binom{a_i}{2} - \sum_j \binom{b_j}{2}\right] - \left[\sum_i \binom{a_i}{2} \sum_j \binom{b_j}{2}\right] / \binom{q}{2}}$,
   where $q_{ij}$ is an element of the contingency table between to the ground-truth and inferred cell attachment matrix, $a_i$ is the sum of the $i$-th row and $b_j$ the sum of the $j$-th column of the contingency table and $q$ is the dimension of the contingency table.
3. *accuracy on corrected genotype*,
   as computed by comparing the corrected (theoretical) genotype matrix with the ground-truth genotype matrix, i.e., prior to the introduction of noise and missing data in the synthetic data generation step.

##### 1.7 Summary of notation

| Symbol | Description |
|---|---|
| $t_i$ | $i$-th time point |
| $m_i$ | Number of variants in the $i$-th experiments (e.g., time point) |
| $\tilde{m}$ | Number of variants among all experiment |
| $y$ | Number of time points |
| $k$ | Number of genotypes in the model |
| $n_i$ | Number of cells in a time point $t_i$ |
| $\alpha_i$ | False Positive Rate of the $i$-th experiment |
| $\beta_i$ | False Negative Rate of the $i$-th experiment |
| $\gamma_i$ | Ratio of missing entries of the $i$-th experiment |
| $\mathbf{G}$ | Estimated genotype matrix |
| $\mathbf{B}$ | Perfect phylogenetic matrix |
| $\mathbf{D}_i$ | Input mutational profile matrix of the $i$-th experiment |
| $\tilde{\mathbf{D}}_i$ | Expanded input data matrix of the $i$-th experiment |
| $\mathbf{C}_i$ | Cell attachment matrix of the $i$-th experiment |
| $\tilde{\mathbf{C}}_i$ | Expanded cell attachment matrix of the $i$-th experiment |
| $\tilde{\mathbf{G}}$ | Estimated expanded genotype matrix |
| $\tilde{\mathbf{G}}^*$ | Corrected (theoretical) expanded genotype matrix |
| $b_{i,j}$ | $i, j$ element of the matrix $\mathbf{B}$ |
| $\mathbf{c}^i_{r,l}$ | The element of the $r$-th row and of the $l$-th column of $\mathbf{C}_i$ |
| $d_{i,j}$ | $i, j$ element of the matrix $\mathbf{D}$ |
| $c_{i,j}$ | $i, j$ element of the matrix $\mathbf{C}$ |
| $\mathbf{b}_i$ | $i$-th row of the phylogenetic matrix $\mathbf{B}$ |
| $w_i$ | Weights of the likelihood function of the $i$-th time point |
| $\rho_{\mathbf{B}',\mathbf{C}'}$ | Ratio of acceptance of a (i.e., the proposal of a $\mathbf{B}'$ and $\mathbf{C}'$ ) in the MCMC |
| $T$ | Learning rate parameter in the MCMC |
| $V$ | Set of vertices of a single-cell longitudinal clonal tree |
| $v_k$ | $k$-th vertex of a single-cell longitudinal clonal tree |
| $E$ | Set of edges of a single-cell longitudinal clonal tree |
| $P$ | Set of weights of a single-cell longitudinal clonal tree |
| $p_k$ | Weight function associated to the $k$-th vertex |
| $\mathcal{E}(i, j)$ | Euclidean distance between two columns of $\tilde{\mathbf{G}}^*$ |
| ARI | Adjusted Rand index |
| TP | True positive |
| FP | False positive |
| TN | True negative |
| FN | False negative |

#### 2 Additional results on simulations

##### 2.1 Cancer population dynamics simulator

To generate synthetic single-cell mutational profiles, we employed the cancer population dynamics simulator presented and used in [4, 5]. Precisely, a stochastic branching process is simulated, in which at every time step (named generation) a cell can replicate itself, with a given replication rate $r_{norm}$, and/or acquire a mutation with probability $p_{mut}$. Every time a new mutation is acquired we have a probability $p_{dri}$ that such mutation is a driver, which provides a fitness advantage $f_{dri}$ to the cell, i.e, an increase in the replication rate.

We performed a large number of independent simulations, by fixing $p_{mut} = 0.6$, and by scanning two values for $p_{dri} = (10^{-3}, 10^{-4})$ and two values for $f_{dri} = (0.01, 0.02)$. For each combination of parameters, we performed 50 simulations up to generation $t = 300$. We finally selected 15 simulations in which a number of drivers between 7 and 15 was observed and we sampled the genotypes of cells at time $t = 50, 150, 300$ to generate the mutational profiles to be used as inputs for LACE.

In particular, we defined three simulated settings, by sampling longitudinal datasets with different number of cells, and by inflating data with false positives $\alpha$, false negatives $\beta$ and missing entries $\gamma$, as shown in Supplementary Table 1.

##### 2.2 Simulation settings

**Population dynamics simulator [4, 5]**

| Simulation runs | Simulation time | $p_{mut}$ | $p_{dri}$ | $f_{dri}$ |
|---|---|---|---|---|
| 50 × setting | 300 generations | 0.6 | $(10^{-4}, 10^{-3})$ | (0.01,0.02) |

**Single cell dataset generation**

| Setting | N. of datasets | Sampling time | $\alpha$ | $\beta$ | $\gamma$ | $n$ |
|---|---|---|---|---|---|---|
| **(A) Distinct plaforms** | 20 × 15 topologies | $t = 50$ | 0.01 | 0.1 | U∼(0,0.1) | 100 |
| | | $t = 150$ | 0.03 | 0.3 | U∼(0,0.1) | 1000 |
| | | $t = 300$ | 0.01 | 0.1 | U∼(0,0.1) | 100 |
| **(B) Tech variability low noise** | 20 × 15 topologies | $t = (50, 150, 300)$ | U∼(0.01, 0.02) | U∼(0.1, 0.2) | U∼(0,0.1) | U∼(100, 300, 600, 1000) |
| **(C) Tech variability high noise** | 20 × 15 topologies | $t = (50, 150, 300)$ | U∼(0.01, 0.02, 0.03) | U∼(0.1, 0.2, 0.3) | U∼(0,0.1) | U∼(100, 300, 600, 1000) |

**Clonal/mutational tree inference**

| Technique | Input $\alpha, \beta$ | MCMC iterations | Restarts | MCMC move prob | Coverage |
|---|---|---|---|---|---|
| LACE | true values | 1000 | 20 | default (0.5, 0.5) | - |
| SCITE | weighted average (true values) | 1000 | 20 | default | - |
| TRaIT | weighted average (true values) | - | - | - | - |
| SiCloneFit | weighted average (true values) | 1000 | 20 | default | - |
| Sifit | weighted average (true values) | 1000 | 20 | default | - |
| CALDER | - | - | - | - | 200x |

Table 1: **Simulation settings.**

##### 2.3 Additional results on comparative assessment with SCITE, TRaIT and SiCloneFit

We here present additional results on the comparative assessment of LACE, SCITE, TRaIT, Sifit and SiCloneFit on the simulated experiments described in the main text and in Supplementary Table 1.

In Fig. 3 we show: $(i)$ the accuracy of the inferred model with respect to the ground-truth topology, $(ii)$ the ARI with respect to the ground-truth cell attachment matrix, $(iii)$ the accuracy of the corrected genotype matrix with respect to the ground-truth genotype matrix.

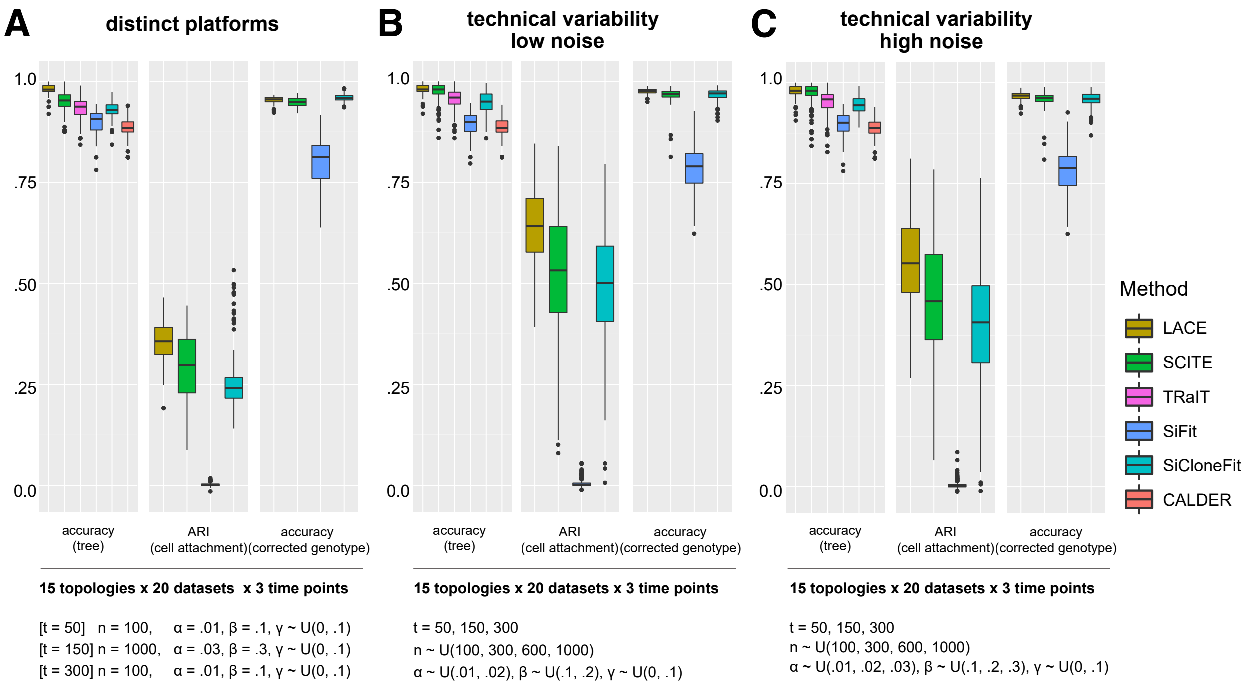

Figure 3: **Comparison on simulated data (further metrics).** We show the results of the comparative assessment described in the main text and in Table 1, in which we compare LACE with SCITE [2], TRaIT [6], Sifit [7], SiCloneFit [8] and CALDER [5]. We here show the distribution of accuracy with respect to the ground-truth topology, of adjusted Rand index with respect to the ground-truth cell attachment matrix and of accuracy with respect to the ground-truth genotype (further metrics are shown in Fig. 2 of the main text). Notice that genotype metrics could not be computed for TRaIT and CALDER, as such methods do not provide cell attachments or corrected genotypes.

In Fig. 4 one can see the scatterplot returning the precision and recall values (with respect to the ground-truth tree topology) for each simulation in all settings, with respect to all tested methods. In the graph, experiments are grouped in bins. LACE proves to outperform all competing methods in all settings.

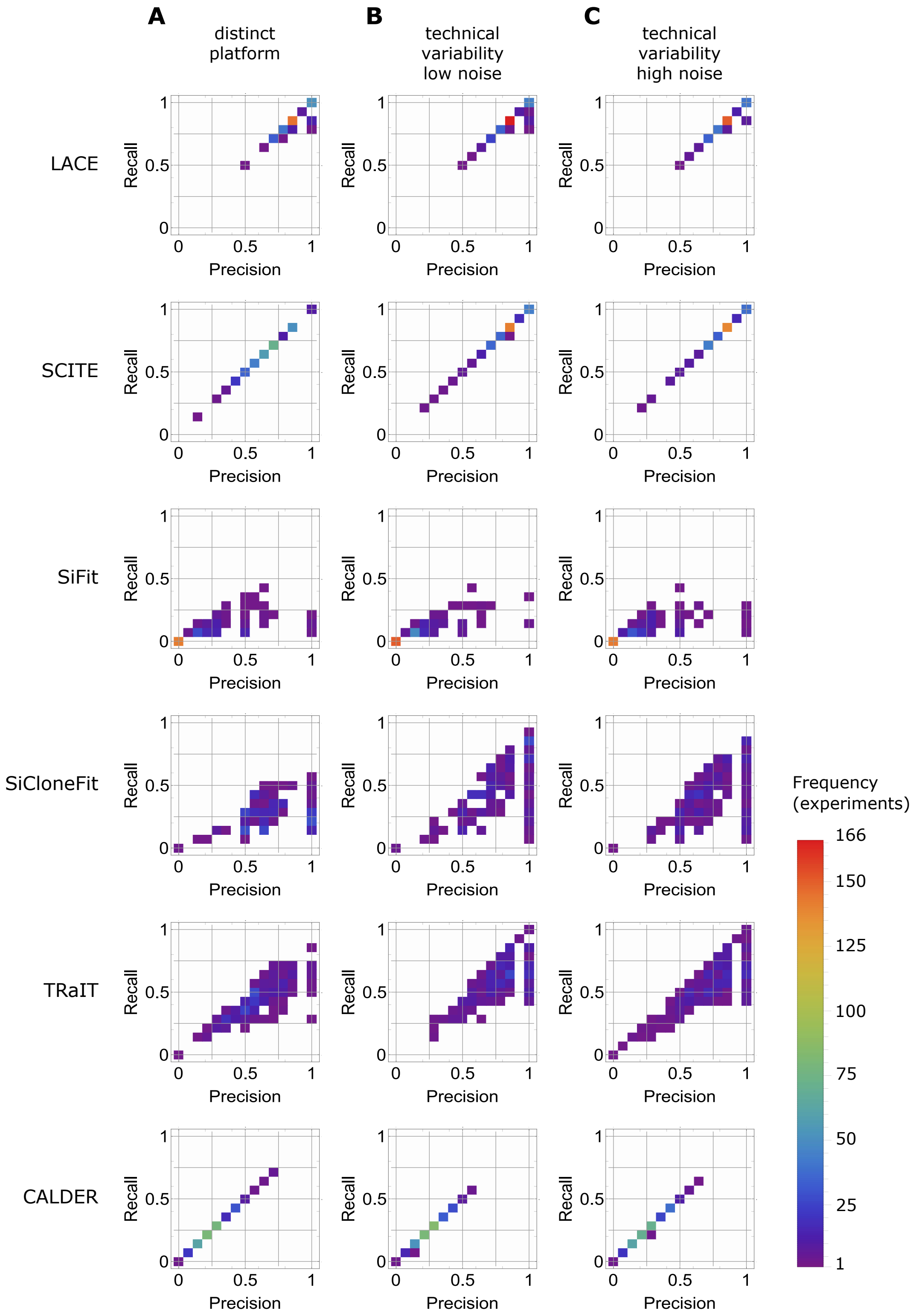

Figure 4: **Precision and recall on simulated data.** The values of precision and recall (with respect to the ground-truth tree topology) for each inference on all simulation settings is displayed as scatter-plot with respect to all considered techniques, namely: LACE, SCITE [2], SiFit [7], SiCloneFit [8], CALDER [5] and TRaIT [6]. In the plot every experiment is represent as a couple $\{x, y\}$ where $x$ =precision and $y$ =recall; every axis is divided into 15 equal bins, for which the observed frequency is returned for every method.

#### 2.4 Tests on simulated data with violations of the Infinite Sites Assumption

We assessed the robustness of the results produced by LACE when data include violations of the Infinite Sites Assumption in terms of back mutations, i.e., the loss of one or more mutations previously acquired during the evolutionary history of the tumor and due to, e.g., loss of heterozygosity.

In detail, we employed 5 binary datasets sampled from each of the 15 topologies of setting A – distinct platforms (see Supplementary Table 1) and we mimicked the loss of a number of mutations, with the following procedure.

By starting from the ground-truth genotype matrix, we first selected a ratio $r$ of variants, among those present in at least 1 single cell at time $t_2$, to be affected by back mutations; for each selected variant, we then picked a ratio $f$ of randomly chosen entries to be flipped to 0 at time $t_3$, among the entries equal to 1 in the ground-truth genotype matrix. We finally inflated the resulting dataset, now including back mutations, with false positives, false negatives and missing data, in rates described in Table 1 for Setting A. Accordingly, we generated the following configurations: *(i)* $r = 25\%$, $f = 25\%$, *(ii)* $r = 25\%$, $f = 50\%$, *(iii)* $r = 50\%$, $f = 25\%$, *(iv)* $r = 50\%$, $f = 50\%$

LACE was executed with 1000 MCMC iterations and 20 restarts. In Supplementary Fig. 5 we show the distribution of precision, recall and accuracy with respect to the ground-truth topology and of the accuracy of the corrected genotype matrix with respect to the ground-truth genotype matrix including back mutations – in the 4 scenarios described above. As one can notice, the performance of LACE is robust also in presence of relatively high fractions of variants affected by back mutations.

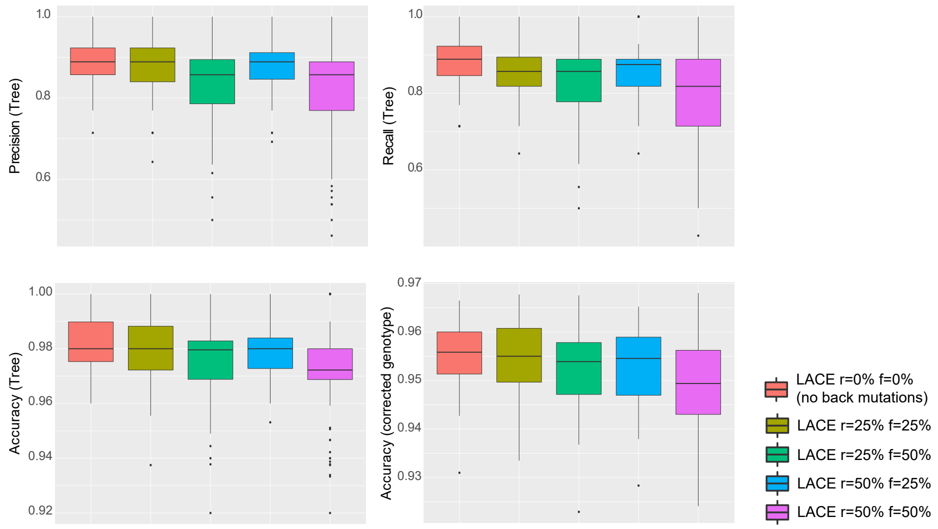

Figure 5: **Performance assessment on simulations with violations of the ISA.**The performance of LACE in distinct scenarios involving violations of the Infinite Sites Assumption is shown. The 4 configurations are described in the text. The box plots display the distribution of accuracy, precision and recall between the ground-truth generative topology and of the accuracy of the corrected genotype matrix with respect to the ground-truth genotype matrix including back mutations. Accuracy and ARI with respect to cell attachament matrix were not computed, as genotypes/clones including back mutations are not modeled in LACE.

##### 2.5 Tests on error rates grid search

We tested on simulations the performance of LACE in case the true values for the false positive rate $\alpha$ and the false negative rate $\beta$ are not provided as input. In this case, LACE allows to employ a built-in grid search over a user-defined range of values for $\alpha$ and $\beta$. In detail, we selected 1 binary dataset sampled from each of the 15 topologies of setting C – technical variability high noise (see Supplementary Table 1), which are characterized by randomly assigned values of $\alpha$ and $\beta$ at distinct time points. We run LACE with 1000 MCMC and 20 restarts, by employing the grid-search defined in Supplementary Table 2.

In Supplementary Fig. 6 we compare the performance of LACE in case the true values of $\alpha$ and $\beta$ are provided as input with the case in which the grid-search is employed. As one can notice, the performance of LACE is extremely robust when employing the grid-search, as precision, recall and accuracy with respect the ground-truth tree topology, as well as the ARI on cell attachment and the corrected genotype accuracy are not affected, whereas only a minor decrease in performance is observed with respect to the accuracy of the cell attachment.

| | $\alpha_1$ | $\alpha_2$ | $\alpha_3$ | $\beta_1$ | $\beta_2$ | $\beta_3$ |
|---|---|---|---|---|---|---|
| #1 | 0.01 | 0.01 | 0.01 | 0.10 | 0.10 | 0.10 |
| #2 | 0.02 | 0.01 | 0.01 | 0.20 | 0.10 | 0.10 |
| #3 | 0.03 | 0.01 | 0.01 | 0.30 | 0.10 | 0.10 |
| #4 | 0.01 | 0.02 | 0.01 | 0.10 | 0.20 | 0.10 |
| #5 | 0.02 | 0.02 | 0.01 | 0.20 | 0.20 | 0.10 |
| #6 | 0.03 | 0.02 | 0.01 | 0.30 | 0.20 | 0.10 |
| #7 | 0.01 | 0.03 | 0.01 | 0.10 | 0.30 | 0.10 |
| #8 | 0.02 | 0.03 | 0.01 | 0.20 | 0.30 | 0.10 |
| #9 | 0.03 | 0.03 | 0.01 | 0.30 | 0.30 | 0.10 |
| #10 | 0.01 | 0.01 | 0.02 | 0.10 | 0.10 | 0.20 |
| #11 | 0.02 | 0.01 | 0.02 | 0.20 | 0.10 | 0.20 |
| #12 | 0.03 | 0.01 | 0.02 | 0.30 | 0.10 | 0.20 |
| #13 | 0.01 | 0.02 | 0.02 | 0.10 | 0.20 | 0.20 |
| #14 | 0.02 | 0.02 | 0.02 | 0.20 | 0.20 | 0.20 |
| #15 | 0.03 | 0.02 | 0.02 | 0.30 | 0.20 | 0.20 |
| #16 | 0.01 | 0.03 | 0.02 | 0.10 | 0.30 | 0.20 |
| #17 | 0.02 | 0.03 | 0.02 | 0.20 | 0.30 | 0.20 |
| #18 | 0.03 | 0.03 | 0.02 | 0.30 | 0.30 | 0.20 |
| #19 | 0.01 | 0.01 | 0.03 | 0.10 | 0.10 | 0.30 |
| #20 | 0.02 | 0.01 | 0.03 | 0.20 | 0.10 | 0.30 |
| #21 | 0.03 | 0.01 | 0.03 | 0.30 | 0.10 | 0.30 |
| #22 | 0.01 | 0.02 | 0.03 | 0.10 | 0.20 | 0.30 |
| #23 | 0.02 | 0.02 | 0.03 | 0.20 | 0.20 | 0.30 |
| #24 | 0.03 | 0.02 | 0.03 | 0.30 | 0.20 | 0.30 |
| #25 | 0.01 | 0.03 | 0.03 | 0.10 | 0.30 | 0.30 |
| #26 | 0.02 | 0.03 | 0.03 | 0.20 | 0.30 | 0.30 |
| #27 | 0.03 | 0.03 | 0.03 | 0.30 | 0.30 | 0.30 |

Table 2: **Grid search values.** Error rates grid search employed in the simulated experiments described in the text. 27 combination of $\alpha_i$ and $\beta_i$ are defined.

To show the robustness of the inference accuracy when wrongly specified values of $\alpha$ and $\beta$ are provided as input to LACE, in Supplementary File 3 we display the performance on all metrics with respect to all the error rates combinations shown in Supplementary Table 2 with respect to 1 binary dataset sampled from each of the 15 topologies of setting C – technical variability high noise (see Supplementary Table 1) . The inference of LACE is proven robust with respect to the considered metrics.

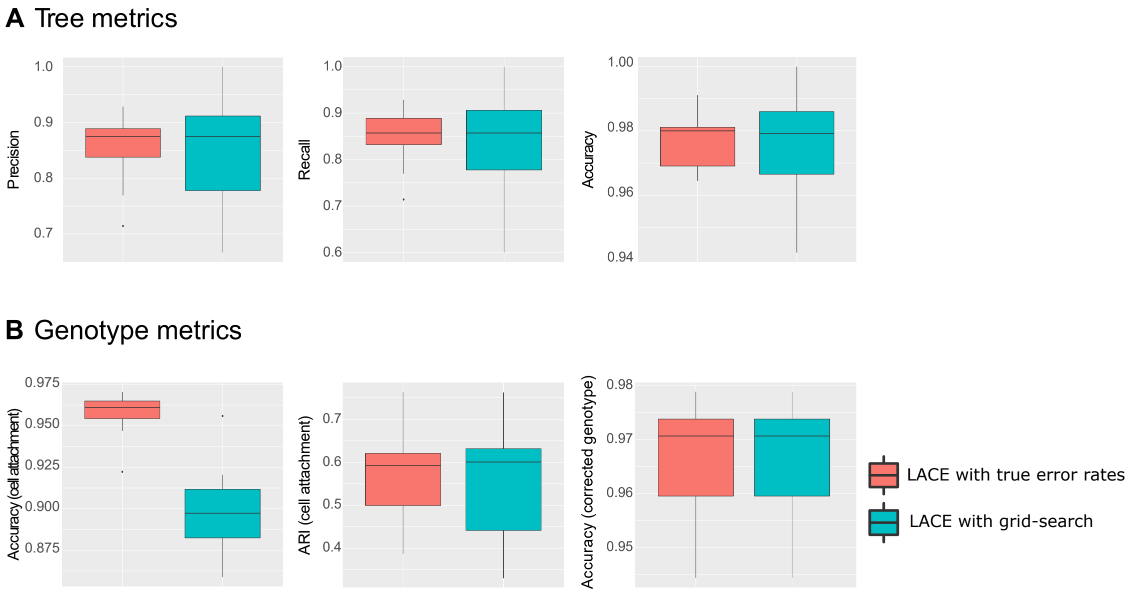

Figure 6: **Performance assessment on error rates grid search.** The comparison of the performance of LACE in case the true error rates are provided as input with that in which the grid-search is employed is shown in terms of standard metrics on simulated data.

#### 3 Additional results on real datasets

##### 3.1 Longitudinal dataset from scRNA-seq data of BRAF-mutant melanoma PDXs

###### 3.1.1 GATK pipeline for variant calling from scRNA-seq data

To generate mutational profiles from the scRNA-seq dataset used in the case study, we employed the GATK Best Practices [9] on the 475 single cells selected after quality check (199 on 674 single cell were discarded due to the high fraction of mitochondrial genes, see Methods).

Namely, we first downloaded the RNA sequences, organized in FASTQ files (one for each single cell), from the GEO dataset with accession number GSE116237 using SRA toolkit. Then, we used Trimmomatic (v. 0.39) to remove the nucleotides with low quality score from the RNA sequences [10]. We then applied the GATK Best Practices to call SNVs and Indels in single cells. In brief, we aligned the sequences on the human reference genome (GRCh38 release) using the STAR aligner in 2-pass mode [11]. Then, we used Picard tools to process the SAM files in order to add read groups, sort, mark duplicates and indexing. Afterwards, we used GATK (v. 3.8.1) to hard clip intronic regions with SplitNCigarReads utility and to re-calibrate base alignment by using BaseRecalibrator utility. The latter step requires the information about known single nucleotide polymorphisms (SNPs), so we used the dbSNP 1000 genome project phase 3 database. Finally, we used HaplotypeCaller and VariantFiltration to call genotype variants and filter out those with low quality score (using default parameters).

After applying the GATK pipeline, we obtain a VCF file for each single cell, which were merged in a unique VCF file for the whole dataset. In the Methods section of the main text we discuss the filters successively employed to define the final list of 6 candidate drivers. In Supplementary Fig. 7 the oncoprint returning the mutational profiles of the 475 single cells of PDX MEL006 is displayed.

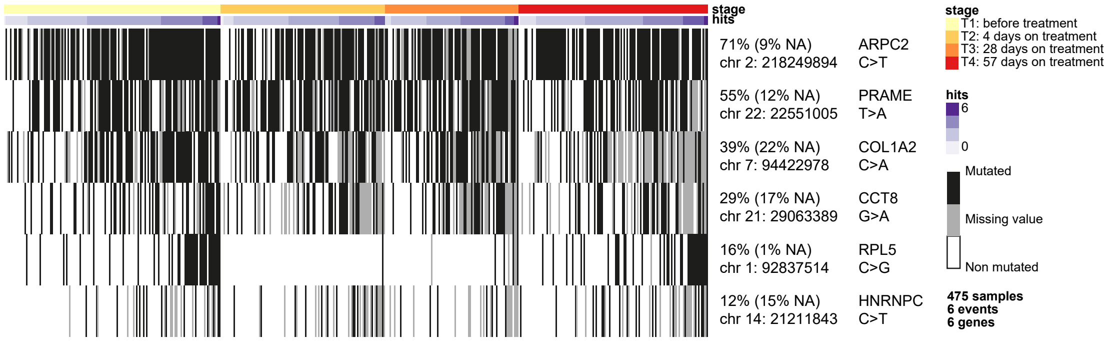

Figure 7: **Oncoprint – PDX MEL006.** Oncoprint returning the mutational profiles of the 475 single cells used in the analysis, from PDX MEL006 [12]. Rows represent single-nucleotide variants, as selected with the procedure described in the text, columns represent single cells, which are grouped for time point. Black cells in the oncoprint represent single cells in which the SNV is present, whereas gray cells represent missing values (i.e., SNVs, either detected or undetected, in a position with coverage lower than 3).

##### 3.1.2 Results of other methods

In Supplementary Figs. 8, 9 and 10 we show the evolution models returned by applying TRaIT, SCITE and SiCloneFit with default parameters to the dataset of PDX MEL006 discussed in the main text. In this case, datasets were concatenated prior to the inference. Notice that the adjacency matrix shown in Fig. 3G of the main text includes the parental relationships retrieved by SiCloneFit that are consistent with perfect phylogenetic assumptions.

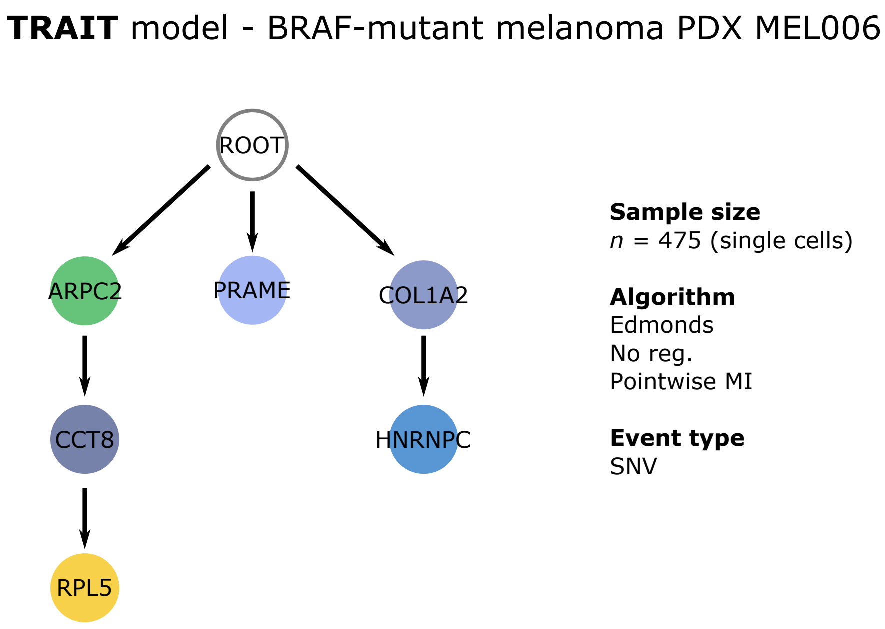

Figure 8: **TRaIT model – PDX MEL006.** The mutational tree of the PDX MEL006 derived from a BRAF-mutant melanoma [12] returned by TRaIT [6] (default parameters) is displayed.

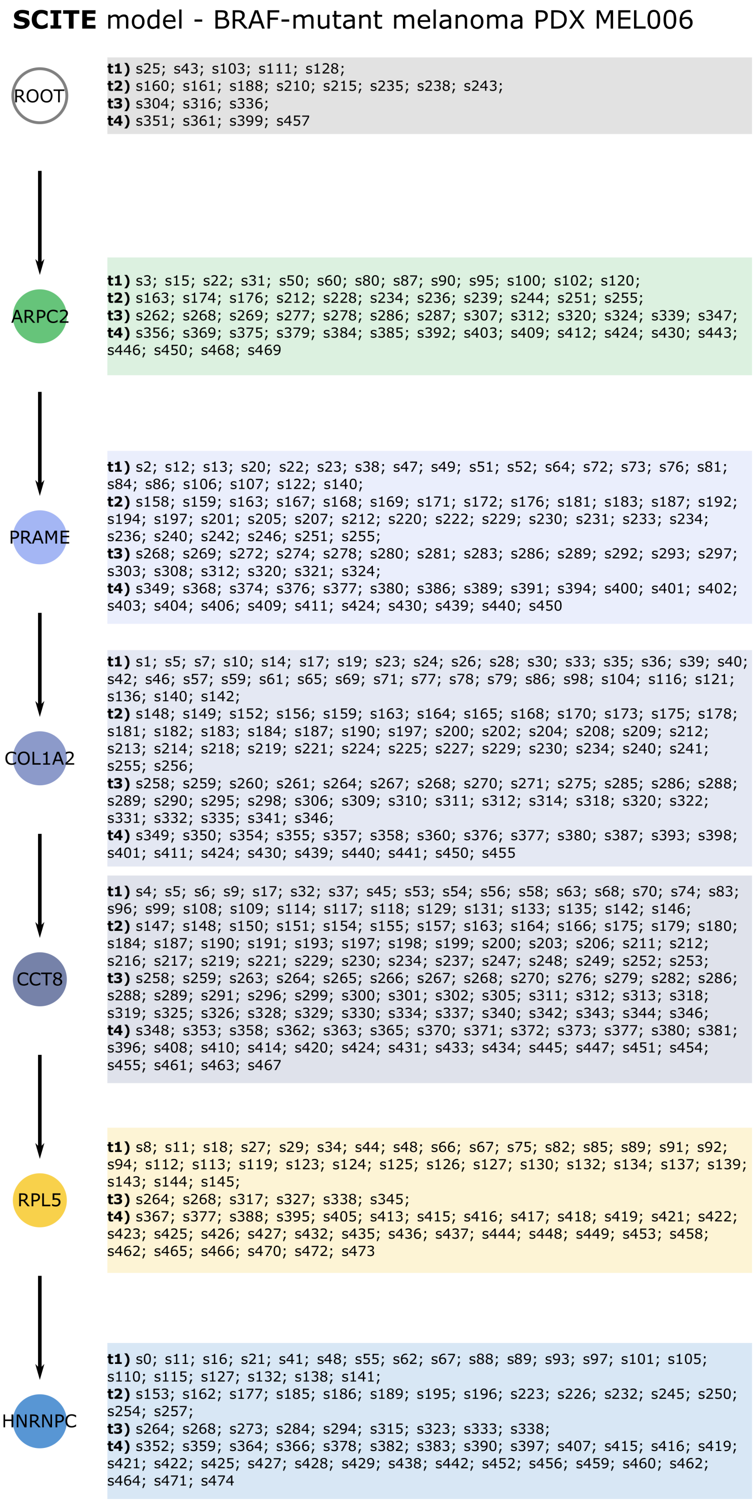

Figure 9: **SCITE model – PDX MEL006.** The mutational tree of the PDX MEL006 derived from a BRAF-mutant melanoma [12] returned by SCITE [2] (default parameters) is displayed. The single cells attached to each node are grouped by time point and ordered as in original dataset.

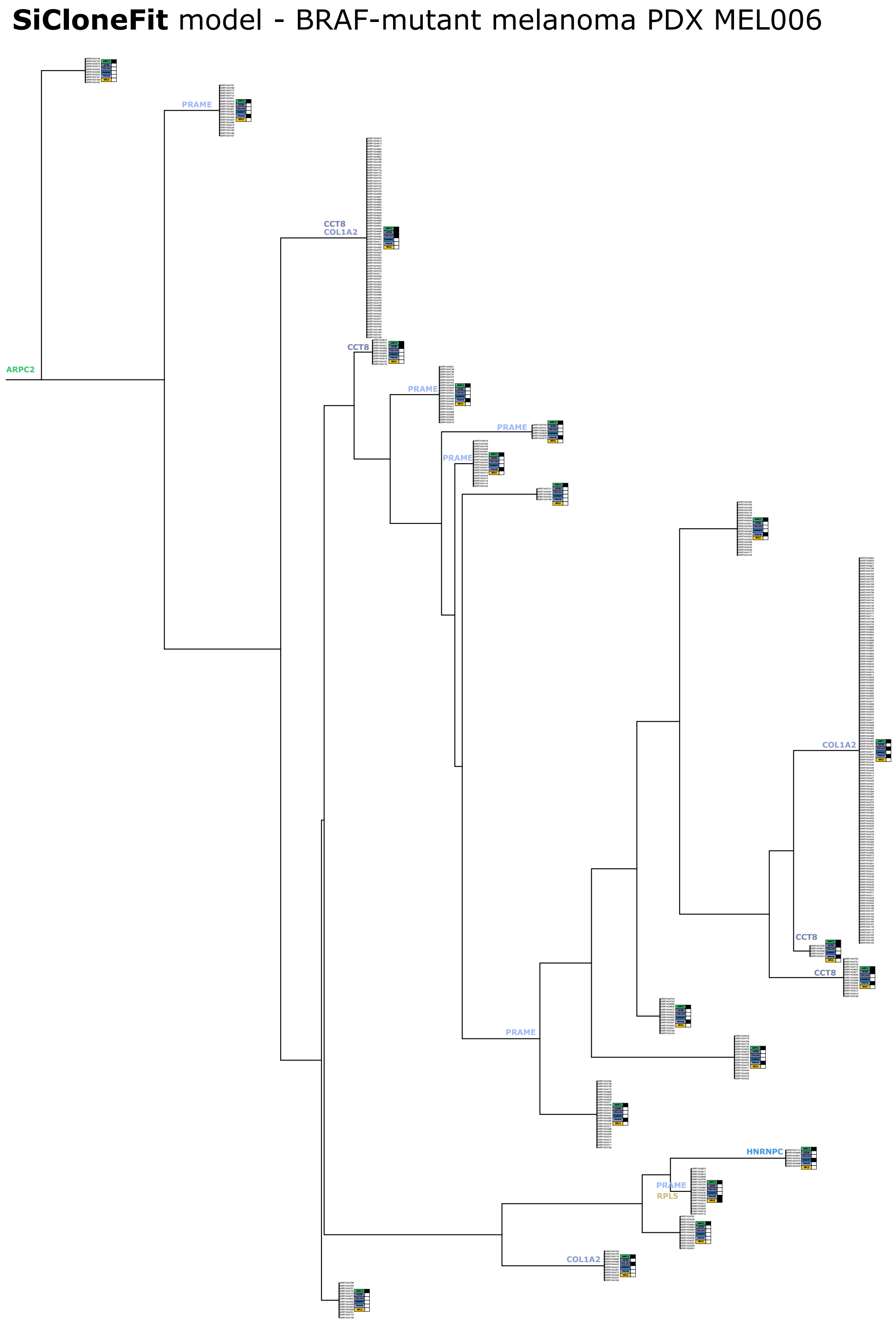

Figure 10: **SiCloneFit model – PDX MEL006.** The phylogenetic tree of the PDX MEL006 derived from a BRAF-mutant melanoma [12] returned by SiCloneFit [8] (default parameters) is displayed. Expected genotypes are displayed near to the inferred clones.

##### 3.1.3 Differential gene expression analysis

We performed differential expression analysis to identify the most significantly up- and down-regulated genes in the distinct (sub)clones identified by LACE on the mutational profiles obtained from PDX MEL006 of [12].

By attaching cells to candidate (sub)clones, LACE allows to partition single cells in distinct groups. In particular, LACE identified a clonal trunk (with mutation on ARPC2) and two subclones, characterized by initiating mutations on RPL5 and PRAME, respectively (see Fig. 3 of the main text).

In order to perform the analysis, we first removed mitochondrial genes, pseudogenes and genes without HUGO symbol; we then normalized the total expression counts of each single cell by library size; ANOVA was performed on $log_2$ transformed data with respect to the 3 partitions of single cells belonging to the distinct subclones.

In particular we first performed the analysis by considering all time points (e.g., the $\text{PRAME}^{\text{MUT}}$ group includes all $\text{PRAME}^{\text{MUT}}$ cells at any time point). In this way it was possible to identify a list of significantly up- and down-regulated genes among the three single-cell partitions. In Supplementary File 1, one can find the list of genes displaying a False Discovery Rate adjusted $p < 0.20$, in addition to the $\log_2$-foldchange as computed on counts normalized by library size.

A second differential expression analysis was performed by considering separately the distinct time points (in this case, for instance, the $\text{PRAME}^{\text{MUT}}$ group includes all $\text{PRAME}^{\text{MUT}}$ cells at a given time point). In this case, no gene appears to be significantly differentially expressed among (sub)clones at time point $t_0$ and $t_3$, whereas a list of 2 and 156 genes are differentially expressed with FDR $p < 0.20$ at time point $t_1$ and $t_2$, respectively. The list of differentially expressed genes with respect to the distinct time points is included in Supplementary File 1.

In Supplementary Fig. 11 one can find the distribution of gene expression values of the 5 genes are significantly up-regulated in $\text{PRAME}^{\text{MUT}}$ cells at time $t_2$ and displaying a $\log_2$-FC larger than 3, namely NGLY1 ($\log_2$-FC $= 4.28$), CDCA7 ($\log_2$-FC $= 3.45$), HK1 ($\log_2$-FC $= 3.27$), DNAJB4 ($\log_2$-FC $= 3.27$), ISOC2 ($\log_2$-FC $= 3.11$).

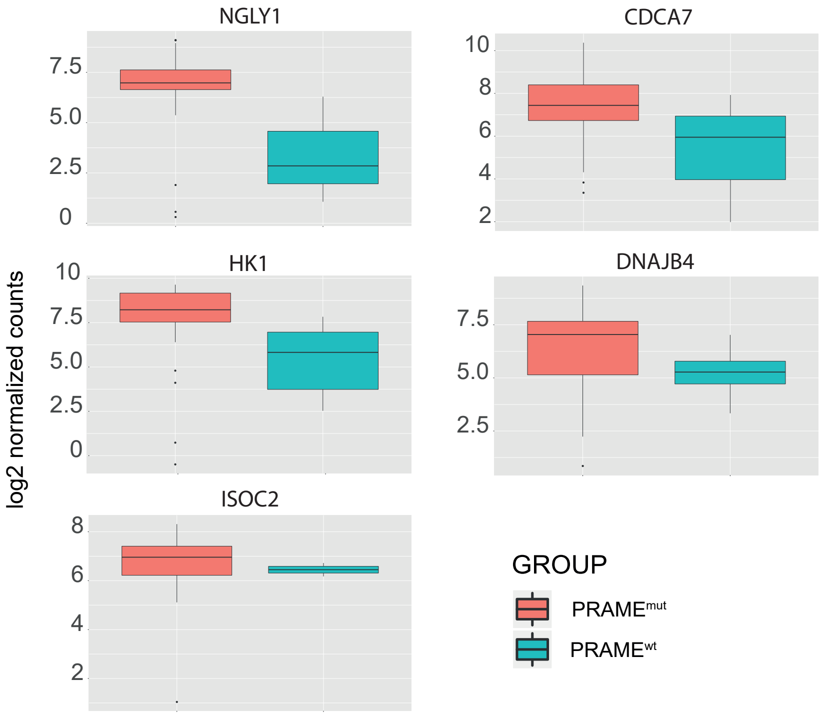

Figure 11: **Gene expression distribution on 5 selected genes – PDX MEL006.** Boxplots representing the $log_2$ normalized count distribution for genes NGLY1, CDCA7, HK1, DNAJB4 and ISOC2, with respect to $\text{PRAME}^{\text{MUT}}$ and $\text{PRAME}^{\text{WT}}$ cells; time point $t_2$ = 28 days.

##### 3.1.4 Single-cell transcriptomic analysis

The single-cell transcriptomic analysis presented in the main text was performed with SCANPY [13] and can be reproduced via the Jupyter notebook included in the online repository available at this link: `https://github.com/BIMIB-DISCo/LACE-UTILITIES`.

In particular, raw expression data were normalized by library size and $\log_2$-transformed. Filtering steps were applied to both samples (i.e., single-cells) and genes, according to the procedure described in the Methods section of the main text. Cells were annotated according to clonal identity (clonal trunk, $PRAME^{MUT}$ and $RPL5^{MUT}$) and cell cycle phase, as estimated via the `score_genes_cell_cycle` SCANPY function on 97 cell cycle genes. Diffusion maps [14] were computed on 58 differentially expressed genes displaying FDR $p < 0.1$ at time $t_2$, by first computing a neighborhood graph of observations (neighborhood = 100, UMAP method). Single cells were finally displayed on the map at different time points.

In Supplementary Fig. 12 we show the diffusion maps for the 475 single cells used in the analysis from PDX MEL006, as computed on 97 cell cycle genes via SCANPY [13].

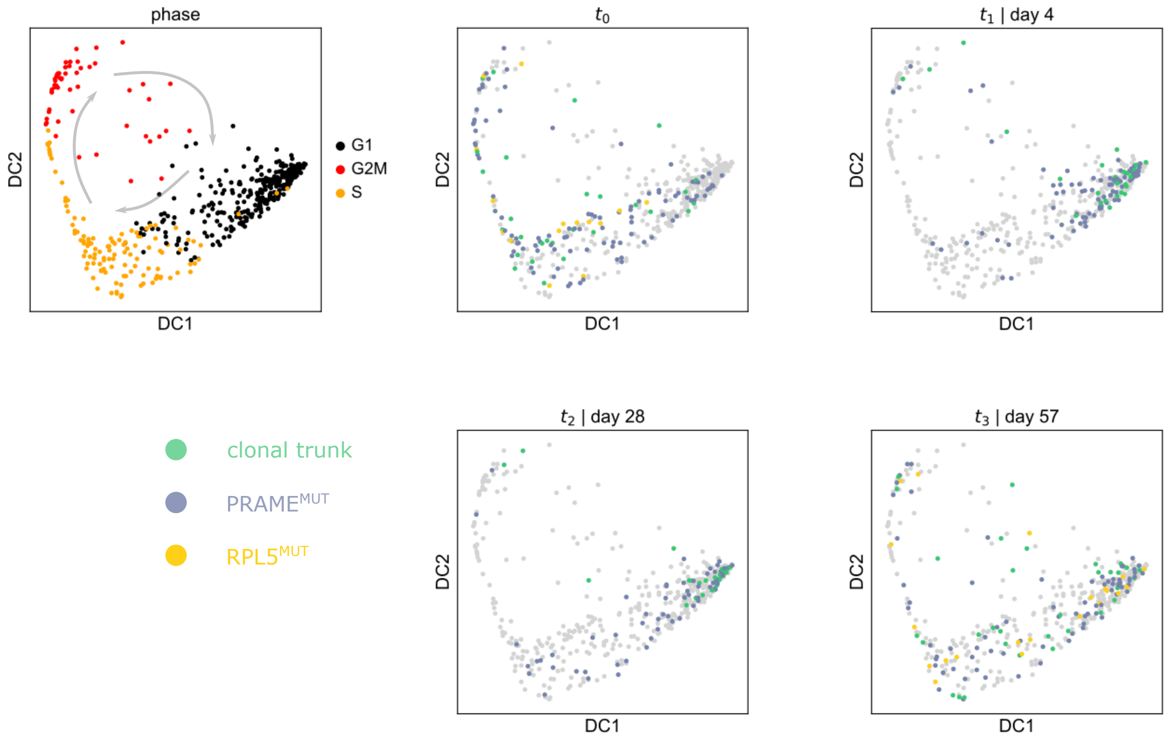

Figure 12: **Diffusion maps on cell cycle genes – PDX MEL006.** Diffusion maps [14] for the 475 single cells used in the analysis from PDX MEL006 [12], as computed on 97 cell cycle genes via SCANPY [13]. The upper-left map shows the cell cycle phase of all single cells at all time points, as estimated via SCANPY, whereas the remaining maps return the clonal identity for all single cells present at distinct time points.

##### 3.1.5 Robustness analysis on variant selection and down/oversampling

We tested the stability of the results delivered by LACE on real data by assessing the impact of: $(i)$ downsampling and oversampling of single cells from the original dataset, with different sample sizes and $(ii)$ selecting either subsets or supersets of of somatic variants provided as input. We used as benchmark the results of the inference presented in the main text on PDX MEL006 [12] ($n = 475$ single cells on all time points, $\tilde{m} = 6$ candidate drivers).

In particular, we defined a first experimental setting, in which we provided as input to LACE datasets with different sample size $n = 333, 380, 428, 523, 570, 618$, which respectively correspond to $-30\%, -20\%, -10\%, +10\%, +20\%, +30\%$ of the cells used in the case study (i.e., $n = 475$). Cells were randomly sampled from the 674 single cells included the original dataset.

In the second setting we executed LACE by providing different numbers of input variants $\tilde{m} = 4, 5, 7, 8$ (the mutational profiles were generated with the procedure described in the main text). In each experiment, such variants were randomly chosen from the list of the following 8 SNVs: ARPC2, CCT8, COL1A2, CYS, HNRNPC,PCBP1, PRAME, RPL5. Such variants are obtained by relaxing the filters on median depth and alternative allele depth, but keeping all the remaining filters described in the main text.

For both experimental settings, LACE was then executed with 10000 MCMC and 50 restarts, for 100 independent runs. In this case we used as ground-truth the topology inferred with the original setting ($n = 475$, $\tilde{m} = 6$) and shown in Fig. 3 of the main text. In Supplementary Figs. 13 and 14 one can see that the results of the inference are extremely stable in presence of under- and over-sampling, as well as when a superset of variants is provided as input, whereas, as expected, a decrease in performance is observed when a small subsets of variants is employed.

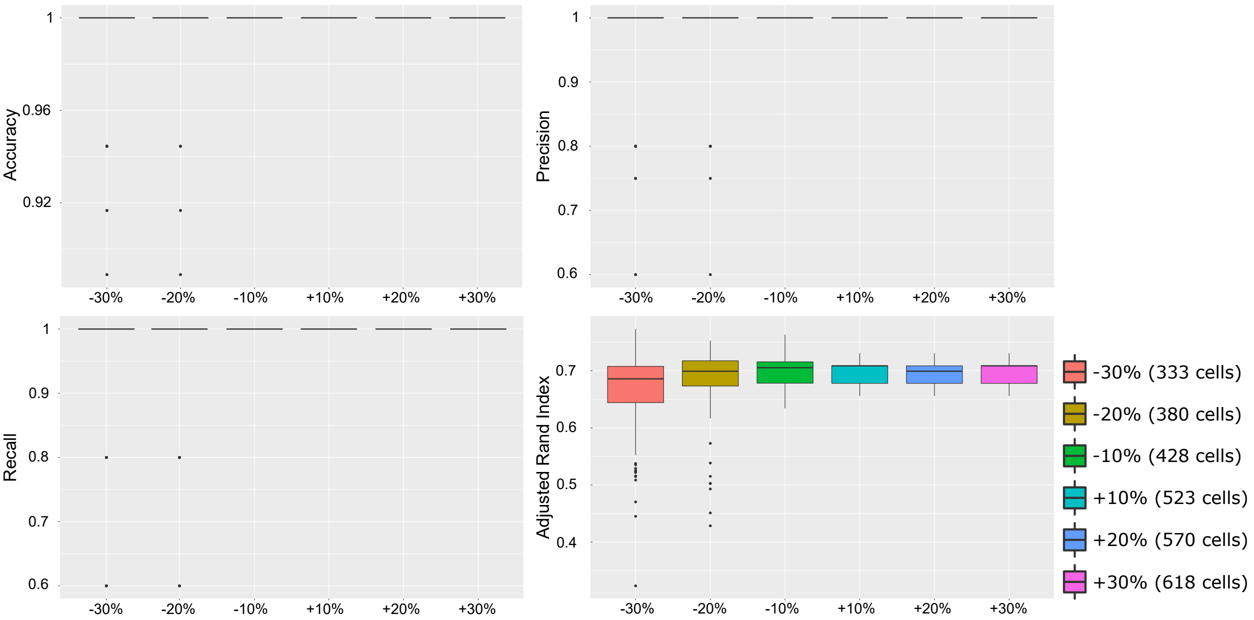

Figure 13: **Robustness on down- and oversampling.** The distribution of accuracy, precision and recall, and the cell attachment adjusted Rand index, as computed with respect to the topology presented in Fig. 3 of the main text, are displayed as boxplots, with regard to the models inferred by LACE when provided with randomly sampled $n = 333, 380, 428, 523, 570, 618$ single cells from the 674 total cells of the original dataset.

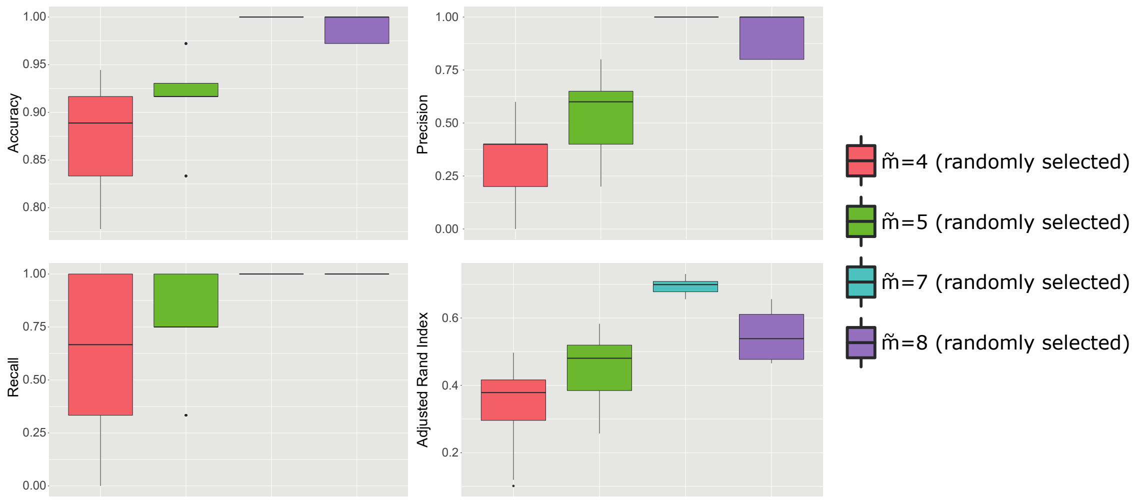

Figure 14: **Robustness on variant selection.** The distribution of accuracy, precision and recall, and the cell attachment adjusted Rand index, as computed with respect to the topology presented in Fig. 3 of the main text, are displayed as boxplots, with regard to the models inferred by LACE when provided with randomly selected $\tilde{m} = 4, 5, 7, 8$ input SNVs from the list presented in the text.

##### 3.2 Longitudinal dataset from targeted scDNA-seq data of breast cancer PDXs

**Application of LACE to longitudinal targeted scDNA-seq dataset from PDXs of ER+ lung metastasis of breast tumors.** We analyzed with LACE a further longitudinal dataset from single-cell targeted DNA-sequencing experiments presented in [15]. Sample SA494 was generated from a lung metastasis of an ER-positive breast tumor and was selected by the authors because of the strong clonal selection after engraftment.

We applied LACE by processing the allelic frequency matrix including a panel of 7 germline and 40 somatic variants selected by the authors, on the metastastic sample (T) and on an engraftment passage (X4) for which single-cell deep re-sequencing was performed. In particular, we excluded germline variants and defined a binary mutational profile, by considering each somatic SNV in each single cell as: $(i)$ present (1) if the allelic frequency $\geq 0.10$, $(ii)$ absent (0) if the allelic frequency is $\leq 0.01$, $(iii)$ missing entry (NA) if the allelic frequency is $> 0.01$ and $< 0.10$ or if already marked as uninformative ($< 25$ mapped reads). Finally, we filtered out all the variants displaying a value of missing data (NA) $\geq 0.15$ in all time points, and variants displaying only 0 or NA values. As a result, the mutational profile matrix provided as input to LACE includes $\tilde{m} = 12$ somatic variants and $n = 42, 56$ single cells for time points $t_0 = \mathsf{T}$ and $t_1 = \mathsf{X4}$, respectively (Supplementary Fig. 15B).

Similarly to the procedure described in the main text, we grouped the variants according to the patterns of co-occurrence across single cells, in order to identify candidate (sub)clones. In this respect, LACE allows to use a similarity measure based on the corrected genotype, which can be employed to cluster the variants of the output model (Supplementary Fig. 15C; see Methods for further details). In Supplementary Fig. 15A the LACE model of sample SA494 is displayed, which offers a picture of the longitudinal evolution of the tumor at the resolution of both genotypes and candidate (sub)clones.

3 candidate (sub)clones are identified by LACE, including a varying number of genotypes, and describing, es expected, a strong clonal selection between the metastatic sample and the mouse xenograft. More in detail, we could identify two subclones emerging from the clonal trunk. Both of them were present in the patient metastasis ($t_0 = \mathsf{T}$), one of which ($\#2$) in extremely low abundance ($\sim 2\%$). After four serial passages of transplanting in mice ($t_1 = \mathsf{X4}$) the rare subclone colonizes the tumor, whereas subclone $\#1$ is completely depleted. Notice that the clonal architecture delivered by LACE is consistent with that proposed by the authors in the original work, yet our method was able to identify a rare clone at the metastatic level, prior to its expansion observed after transplanting (the cell attachment inferred by our analysis and that returned in the original work are provided in Supplementary File 2).

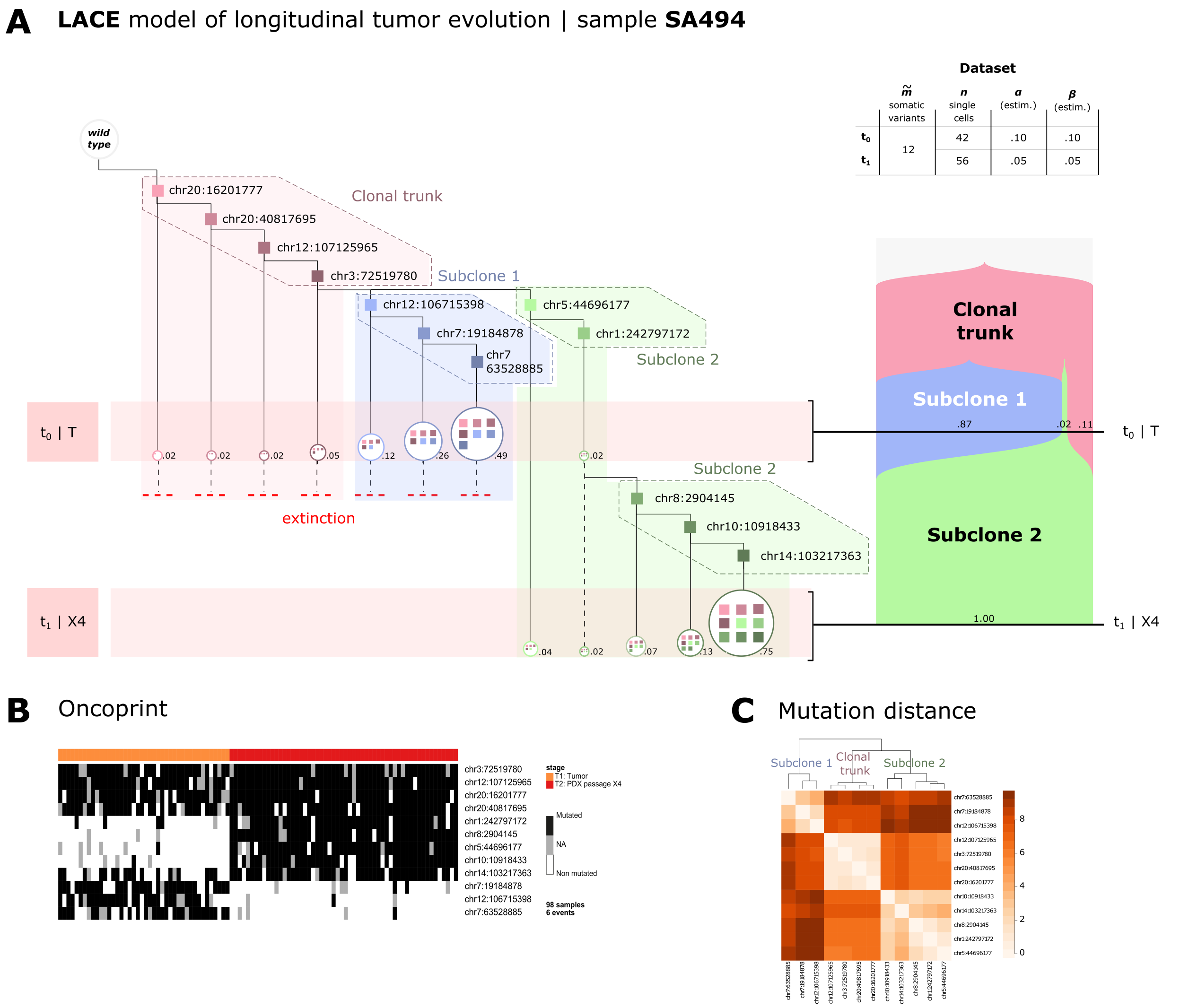

Figure 15: **LACE model – Sample SA494.** **(A)** The longitudinal evolution inferred by LACE from a lung metastasis of a ER-positive breast tumor sample and the corresponding PDX (sample SA494) [15] is displayed. Targeted DNA sequencing experiments were performed on single cells of the lung metastasis (T, $n = 42$ single cells) and of a PDX after four serial passages (X4, $n = 56$), by selecting a panel of 7 germline SNVs and 40 somatic SNVs. Single-cell mutational profiles were generated by discretizing the alternative allele ratio as proposed in the original work. Somatic SNVs displaying a ratio of missing data $> 0.15$ and germline mutations were filtered-out prior to the inference. As a result, $\widetilde{m} = 12$ somatic variants were selected to be used as input in LACE (see the caption of Fig. 4 of the main text for further details). The representation via a standard fishplot, generated via TimeScape [16] at the resolution of candidate (sub)clones, is displayed on the right. **(B)** The oncoprint returning the single-cell mutational profile used in the analysis is shown. **(C)** A heatmap returning the mutational distance among the 12 somatic variants used in the analysis is displayed. 3 clusters are identified which represent candidate (sub)clones and correspond to the 3 genotypes discussed in the original work.
